# Distinct input-output relationships between basal forebrain subnuclei underlie heterogeneous cholinergic modulation of the mouse cortex

**DOI:** 10.64898/2026.11.10.765460

**Authors:** Junhua Tan, Aspen M. Harter, Miranda Kolb, Ryan Lu, Lucas Pinto

## Abstract

The basal forebrain (BF) supports many behavioral functions via its brain-wide projections, which can be recruited independently and task specifically. However, the anatomical underpinnings for this functional heterogeneity are poorly understood. Here, we compared two mouse BF subnuclei, the horizontal diagonal band (HDB) and the substantial innominata (SI), in terms of their anatomical connectivity and functional recruitment. Using anterograde and retrograde tracing, we show that they target largely segregated forebrain regions, but their input comes from a partially overlapping subset of regions. However, many neurons within each region project to just one of the subnuclei, suggesting they can be recruited independently. To test this, we combined optogenetics and widefield Ca^2+^ imaging and showed that stimulating prefrontal cortical projections to the HDB or SI results in distinct large-scale patterns of cortical activity via a cholinergic mechanism. Our results provide an anatomical basis for the flexible, area-specific cholinergic modulation of brain function.

## Introduction

The basal forebrain (BF) is a key neuromodulatory nucleus containing several cell types, among them the cholinergic neurons that provide the only long-range source of acetylcholine (ACh) to the cortex and other forebrain regions^1–4^. These neurons modulate many behavioral processes from sub-second to circadian timescales, including arousal and behavioral state^5–12^, plasticity, learning and memory^13–17^, reward expectation and reinforcement^18–22^, sensory processing, attention and decision making^11,12,23–33^. This functional versatility is thought to be at least in part enabled by a rough topographical arrangement of connections to and from the cortex and elsewhere^34–46^, arising from several subnuclei that comprise the BF^2,47,48^. Indeed, previous studies have revealed highly spatiotemporally heterogeneous patterns of cholinergic release in the cortex, in a task- and sensory-modality specific fashion^6,19,26,27,49^.

However, the anatomical arrangement leading to this functional heterogeneity remains poorly understood. A possibility is that different BF subnuclei receive different input patterns, which would enable their independent recruitment and, in turn, influence distinct sets of downstream regions. To our knowledge, this has not been tested, as no studies have directly bridged the specific anatomical connectivity of BF subnuclei to their function. Moreover, while most recent functional studies have been done in mice, our understanding of BF topography has come primarily from rats and monkeys — most mouse anatomy work has focused on individual subnuclei or subsets of BF targets, and/or considered the BF as a whole^37,50–55^.

To address this gap, here we combined anterograde and retrograde tracing to systematically compare the input-output relationships between two mouse BF subnuclei — the rostroventrally located horizontal diagonal band of Broca (HDB) and the more dorsocaudal subnucleus substantia innominata (SI). We find that they preferentially target largely non-overlapping regions, including in the cortex, and that they receive input from partly non-overlapping neurons from within similar regions. Using optogenetics and widefield imaging, we show that this arrangement is sufficient to generate distinct patterns of large-scale cholinergic modulation of excitatory neurons in the dorsal cortex. Our results establish an anatomical basis for the multiple contributions of BF projections to behavioral function.

## Results

### Basal forebrain subnuclei target distinct forebrain regions

To characterize the output patterns of BF cholinergic neurons from HDB and SI, we analyzed anterograde tracing data from the Allen Mouse Connectivity Atlas^56,57^ (**Fig. 1A**, n = 4 ChAT-IRES-Cre mice for SI and n = 3 for HDB, see STAR Methods for data selection criteria). We co-registered experiments in the Allen Common Coordinate Framework space^58^ (CCFv3) and compared the spatial distributions of cholinergic axons arising from the two subnuclei. We only considered ipsilateral projections, given that contralateral projections were very rare and could have been due to the spread of viral injections to the contralateral hemisphere, particularly for the more medially located HDB (not shown). We first visualized projection density across the forebrain volume (**Fig. 1B**) and projected onto the three major neuroanatomical planes and axes (**Fig. 1C, D**; density is defined as the proportion of projection-containing pixels in a voxel). Compared to SI, HDB preferentially projected more to the medial, caudal and dorsal parts of the cerebrum (**Fig. 1D**, p < 0.001, F-test for nested models). These projection patterns were highly consistent across animals (**Fig. S1**) and generally similar when only considering cortical areas (**Fig. S2**).

**Fig. 1.**
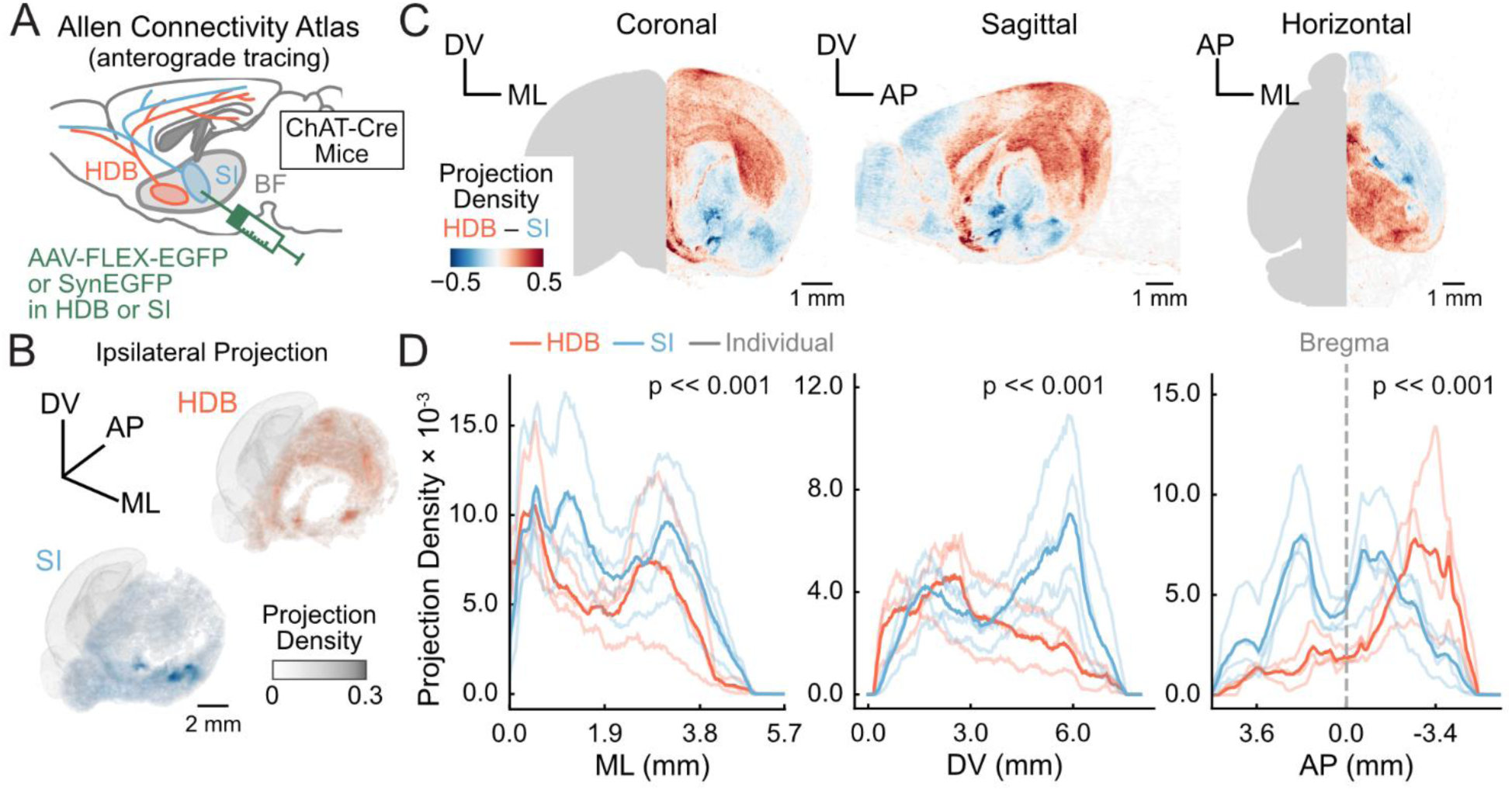
Basal forebrain subnuclei have distinct large-scale projection patterns. (A) Experiment schematics for the publicly available dataset. (B) 3-D reconstruction of average HDB (red) and SI (blue) projection density across the forebrain (n = 3 mice for HDB, n = 4 for SI). AP: anteroposterior; ML: mediolateral; DV: dorsoventral. (C) Average difference between the cholinergic-projection density of the two subnuclei, maximum-projected onto the three neuroanatomical planes. (D) 1-D spatial profiles of projection density from each subnucleus, for individual mice and their averages. The printed p-values are from an F-test for nested linear-regression models with and without interaction terms between injection location and projection density (see STAR methods). For ML, the change in R^2^ between full and reduced models was 0.024, F_df=3_ = 41.389. For DV, the change in R^2^ between full and reduced models was 0.187, F_df=3_ = 338.295. For AP, the change in R^2^ between full and reduced models was 0.168, F_df=3_ = 656.385.

Next, we quantified the distribution of projection strength from each subnucleus to different anatomical regions of the cerebrum (**Fig. 2A**, **S3** and **Table S1**, as defined in the CCFv3 ontology). For each region, we computed the average projection density across voxels, separately for each mouse. The area-level distribution of projection density was significantly different between the subnuclei (**Fig. 2B**, p = 9.47 × 10^-41^, linear regression, Wald test for area × subnucleus interaction; see **Table S1** for all brain regions). Specifically, regions such as the prelimbic cortex (PL), lateral orbitofrontal cortex (ORBl), ventral agranular insular cortex (AIv), basolateral amygdala (BLA) received primarily cholinergic projections from SI (p < 0.05, linear regression, t-test on coefficient contrast). Conversely, the ventral anterior cingulate cortex (ACAv) and retrosplenial (RSPagl, RSPv) cortices, the secondary visual area VISam, and regions of the hippocampal formation (e.g., CA1–3, subiculum SUB) received most input from HDB (p < 0.05).

**Fig. 2.**
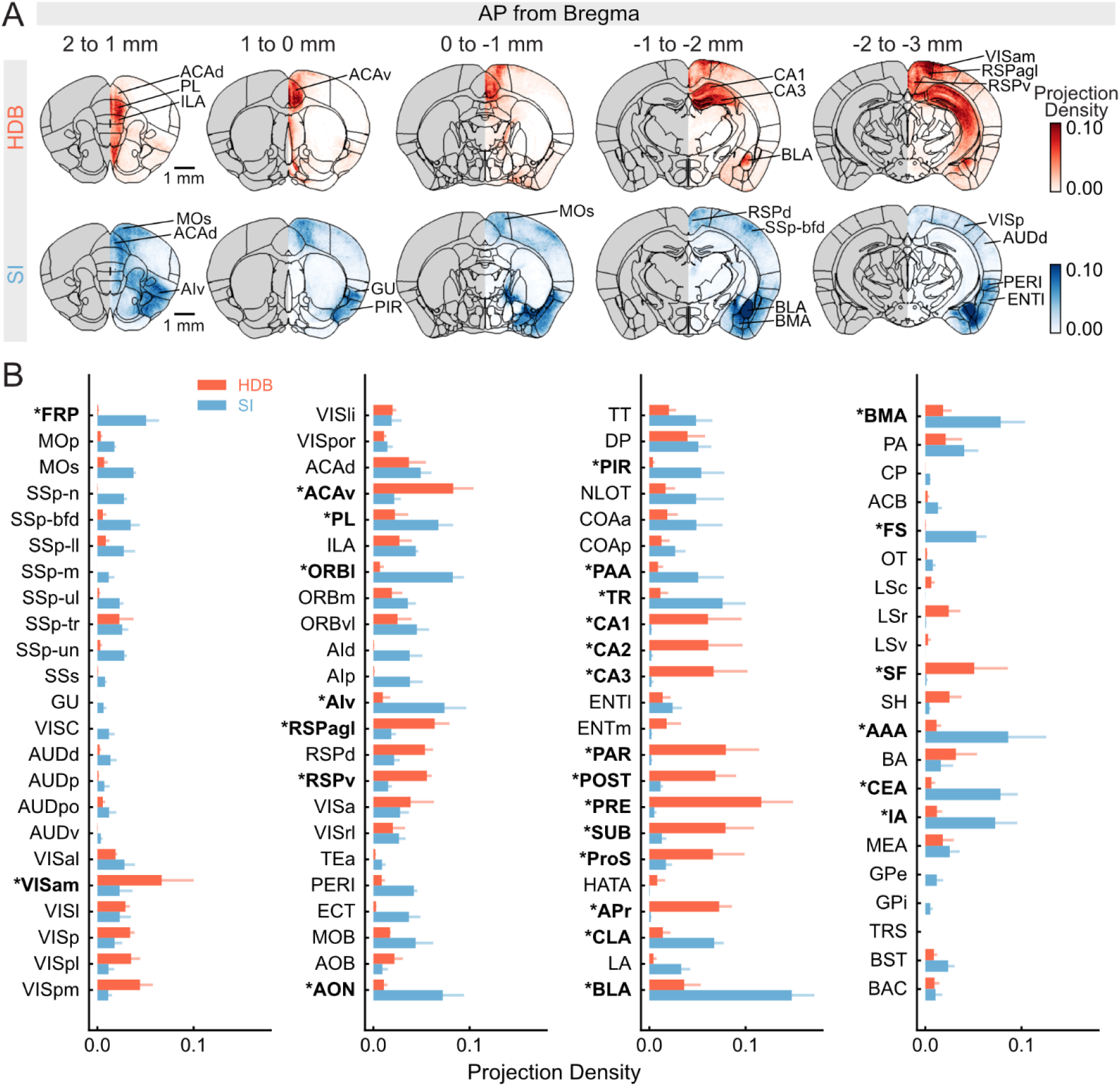
Projection density from HDB and SI to anatomically defined regions of the cerebrum. (A) Cross-mouse average of HDB (red) and SI (blue) projections in coronal view per 1-mm volume (n = 3 for HDB and 4 for SI). (B) Average density of HDB and SI projections to forebrain regions. Error bars, ± S.E.M across mice (n = 3 for HDB and 4 for SI). * and bolded area labels: p < 0.05. This was calculated for each region from a linear regression model with cerebrum regions, injection location (HDB or SI), and their interactions. Coefficient contrast was then calculated for each region as the model-predicted difference between injection locations, and p-values were computed from a *t* distribution and corrected for false discovery rate (FDR). See **Table S1** for area acronym definitions.

Note, however, that although most regions received input predominantly from one subnucleus, several received a mixture of HDB and SI input. Interestingly, a recent study of just the somatosensory cortex showed laminar biases in cholinergic projections, with SI and HDB preferentially targeting deep and superficial cortical layers, respectively^59^. We thus sought to determine whether and to which extent this trend held across the cortex. On aggregate, HDB projections did bias towards superficial layers (and vice-versa for SI, p = 0.001, interaction term of a mixed two-way ANOVA with factors injection location and cortical layer), though with variable laminar patterns across cortical regions (**Fig. S4**). This variability is compatible with a previous study in rats^42^.

Overall, our analysis shows subnucleus-specific cholinergic projection patterns across the cerebrum, indicating that recent findings of large-scale functional heterogeneity could be related to the differential recruitment of BF subnuclei.

### BF subnuclei receive mostly segregated input from a similar subset of regions

Next, we sought to determine whether HDB and SI receive anatomical input from the cerebrum that is segregated enough to enable their differential recruitment during behavior. We performed retrograde AAV tracing, delivering a green and a red fluorophore to each subnucleus (**Figs. 3A, B, S5**, EGFP and mCherry, counterbalanced across mice, n = 4 HDB and 6 SI wild-type mice; 3 of these with both subnuclei injected). Note that this viral strategy is not cell-type specific, meaning that we cannot guarantee that we are mapping input specifically to cholinergic neurons. However, previous rabies-tracing work has shown that glutamatergic, GABAergic and cholinergic neurons in the BF all tend to be targeted by similar brain regions^52^, ameliorating this potential concern.

**Fig. 3.**
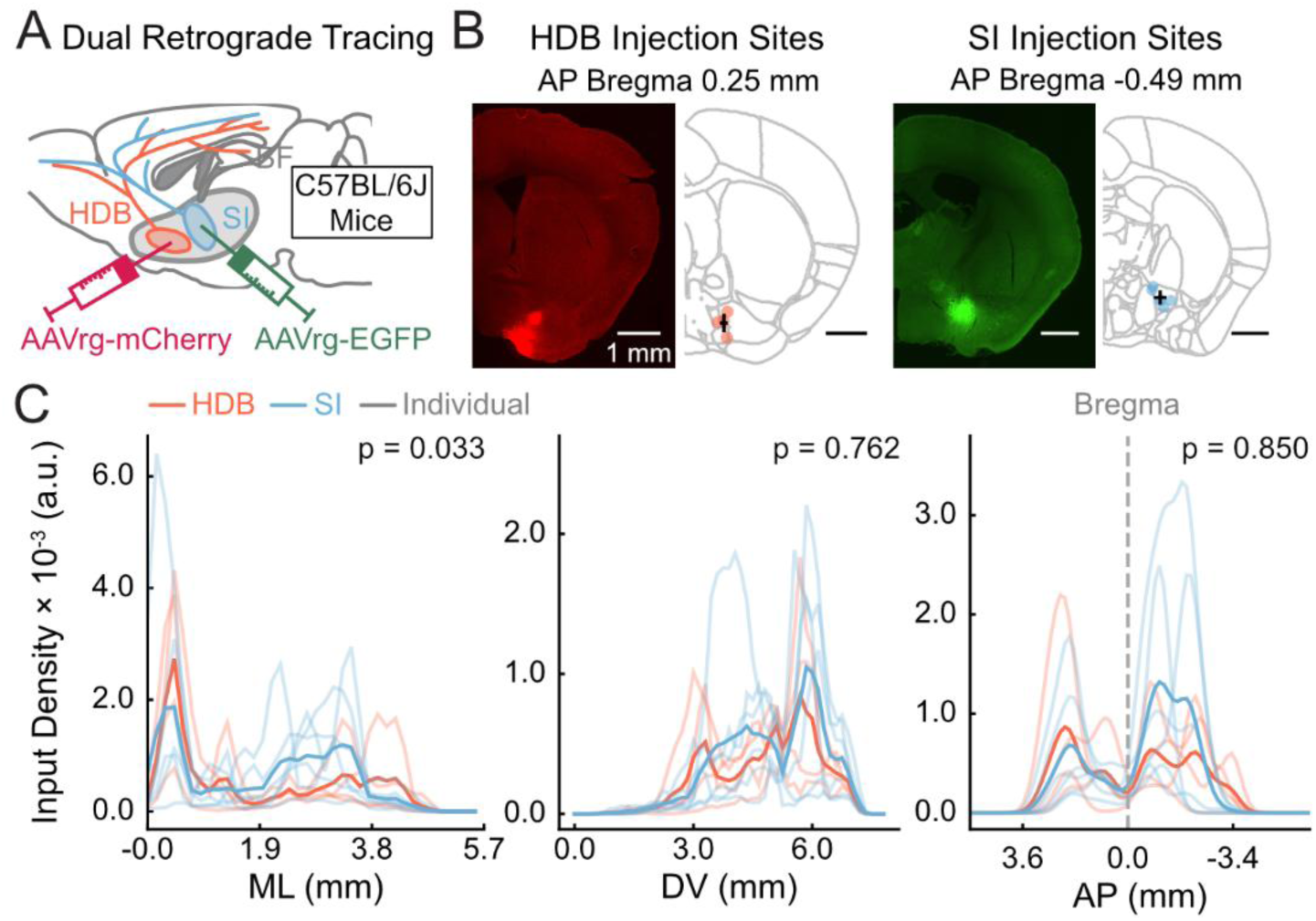
HDB and SI receive a similar overall spatial profile of input from the cerebrum. (A) Dual-color retrograde experiment schematics. (B) Left, example fluorescence images of the HDB (n = 4 mice) and SI (n = 6 mice) injections. Three mice received injections in both subnuclei. Right, centroids of all HDB and SI injection sites used in this study (circles). Crosses are the average and standard deviation across injections. (C) 1-D spatial profiles of input density to each subnucleus along different anatomical axes. The printed p-values are from F-tests as in Fig. 1. For ML, the change in R^2^ between full and reduced models is 0.015, F_df=3_ = 2.947. For DV, the change in R^2^ between full and reduced models is -0.002, F_df=3_ = 0.387. For AP, the change in R^2^ between full and reduced models is -0.002, F_df=3_ = 0.266.

We imaged coronal slices throughout the cerebrum using single-photon epifluorescence microscopy and used the semi-automated QUINT workflow^60^ to co-register our slices to the Allen CCFv3 and segment puncta across the cerebrum expressing either fluorophore (see STAR Methods). We then computed input density as the puncta counts per voxel normalized by the largest puncta count within an experiment. Interestingly, despite their distinct output patterns, we found that input to the HDB and SI was distributed similarly along the DV and AP anatomical axes (**Fig. 3C**, p > 0.05, F-test for nested models), and shifted significantly laterally for SI (p = 0.03). Moreover, although input from the cortex to both subnuclei came predominantly from layer 5 (albeit with area-to-area variability), laminar distributions varied significantly between the subnuclei, being slightly more superficial for HDB (**Fig. S6,** p = 7.46 × 10^-4^, interaction term of a two-way ANOVA with factors injection location and cortical layer).

To further quantify these trends, we compared input density from anatomically defined cerebrum regions to SI or HDB. Compared to their output, input to each subnucleus came from a much smaller subset of regions, comprising mostly the prefrontal cortex (MOs, PL, ILA, ACAd, ACAv), the orbitofrontal cortex (ORBl, ORBm, ORBvl), the insular cortex (AId, AIv, AIp), the pyriform cortex (PIR), and the amygdala (BLA, BMA, CEA) (**Fig. 4A, B**; see **Table S1** for more brain regions). Most of these regions projected to both SI and HDB in similar proportions, but the overall area-level distribution was statistically different between the subnuclei (**Fig. 4B**, p = 0.006, linear regression, Wald test for area × subnucleus interaction). However, only two individual regions had statistically significant differences: PIR projected more to HDB, while the basolateral amygdala (BLA) projected predominantly to SI (p < 0.05, linear regression, t-test on coefficient contrast; though note that the BLA also projected strongly to HDB).

**Fig. 4.**
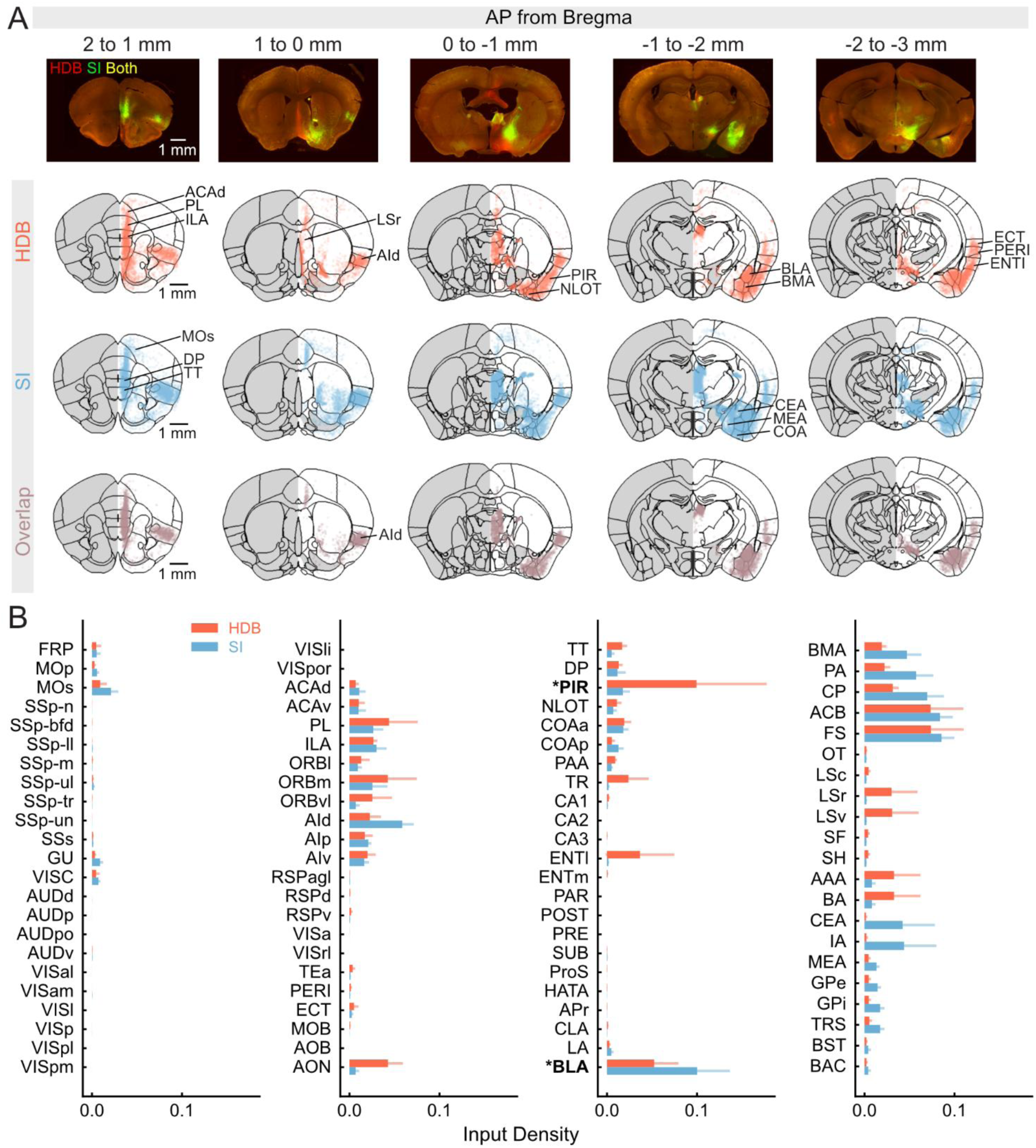
Distributions of input from cerebrum areas to the HDB and SI. (A) Top, representative coronal sections with retrogradely labeled neurons projecting to HDB (red, mCherry) and SI (green, EGFP). Bottom, average of HDB (red) and SI (blue) input puncta, and their spatial overlap, within different 1-mm coronal volumes (n = 4 for HDB and 6 for SI). (B) Average input density to HDB and SI within cerebrum regions. Error bar, ± S.E.M across mice (n = 4 for HDB and 6 for SI). * and bolded area labels: p < 0.05. This was calculated as in Fig. 2. See **Table S1** for area acronym definitions.

The analysis above, however, only allowed us to characterize the overall spatial and areal pattern of input to HDB and SI. Although largely overlapping on these larger spatial scales, this input could still be segregated if subsets of neurons within the same area targeted only one BF subnucleus. To get at this, we used confocal microscopy to measure the amount of within-neuron retrograde tracer overlap in mice with both HDB and SI injections (n = 3 mice, 41 fields of view — FOVs), within six selected cerebrum regions that projected to both subnuclei (**Fig. 5A**) — the dorsal anterior cingulate area (ACAd), prelimbic area (PL), infralimbic area (ILA), dorsal anterior insula (AId), basolateral amygdala (BLA), and cortical amygdala (COA). Across regions, neurons projecting to both subnuclei comprised ∼25–55% of all neurons projecting to HDB and/or SI (**Fig. 5B**, out of a ∼80% overlap ceiling estimated with dual injections in the same location). Thus, in all areas, many neurons project to only one of the BF subnuclei.

**Fig. 5.**
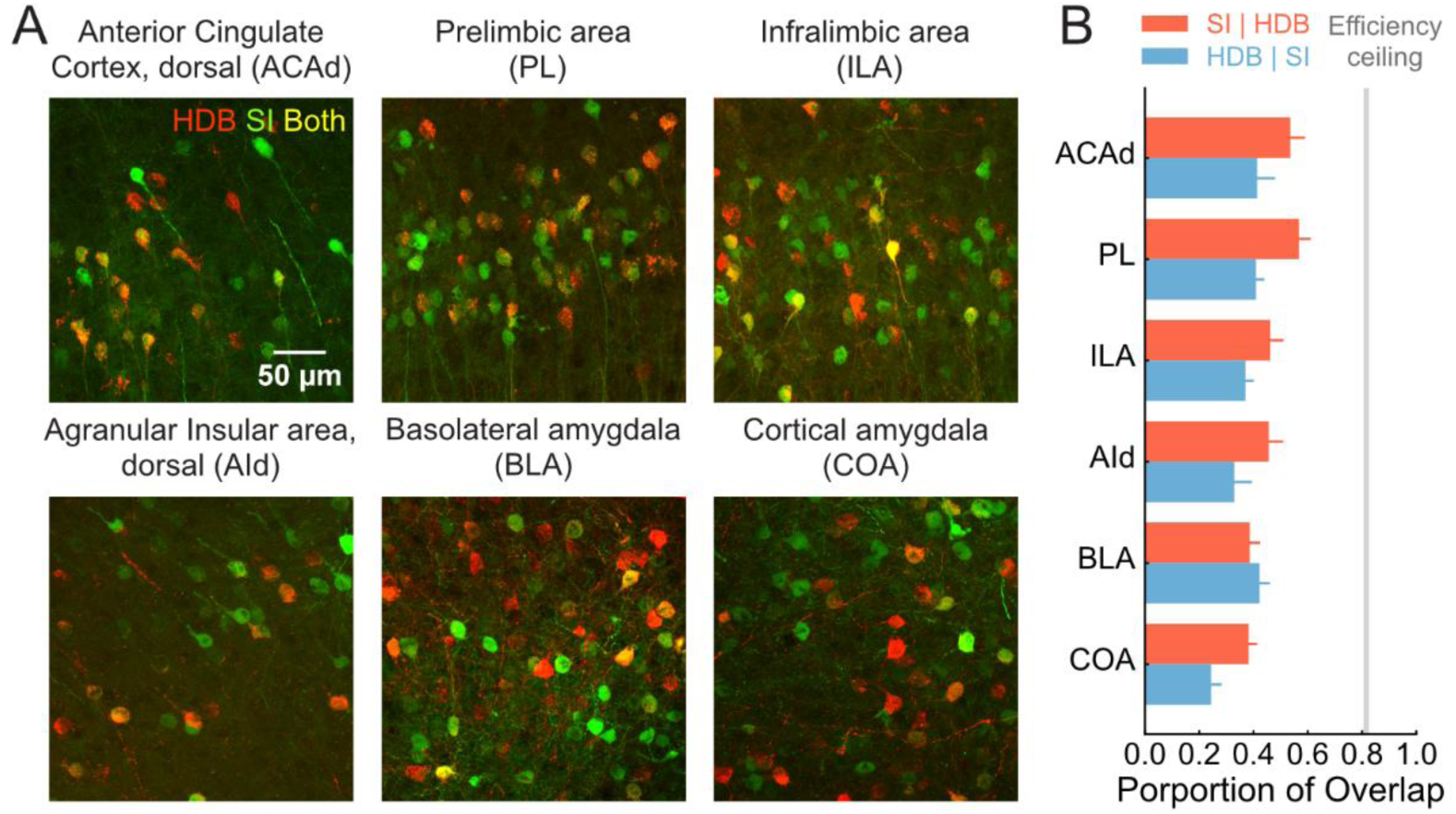
Single-neuron overlap of cortical input to the HDB and SI. (A) Example maximum projection of confocal z-stacks from selected cerebrum regions, with retrogradely labeled neurons projecting to the HDB (red, mCherry), SI (green, EGFP), or both (yellow). (B) Quantification of the proportion overlap for the regions in A. Error bars: ± S.E.M across FOVs (n = 9 for ACAd, PL, and ILA each, n = 4 for AId, n = 6 for BLA, and n = 5 for COA). Gray shaded line: mean ± S.E.M. for the estimate of dual-labeling efficiency ceiling from one mouse injected with both mCherry and EGFP in HDB (n = 208 PL and BLA neurons).

Overall, our retrograde tracing experiments show that, while BF subnuclei receive input from similar forebrain regions (with a few exceptions), the neuronal populations targeting each subnucleus are only partially overlapping. This could provide enough anatomical specificity for differential functional recruitment of these subnuclei, in turn resulting in area-specific patterns of modulation by the BF.

### mPFC→HDB projections indirectly modulate dorsal cortical activity via a cholinergic mechanism

Whether our anatomical results mean that BF subnuclei can be recruited differentially depends on many factors, e.g., how interconnected the neurons targeting them are and how densely HDB and SI connect to each other. Moreover, given that our retrograde tracing strategy lacks cell-type specificity, and that the BF contains multiple cortically projecting cell types^3,61^, it is unclear whether projections to BF subnuclei recruit its cholinergic neurons. To directly answer these questions, we developed an experimental paradigm to simultaneously stimulate input to the BF with optogenetics and record activity from across the dorsal cortex with mesoscale widefield Ca^2+^ imaging, of either cholinergic axons or cortical excitatory neurons. We selected one of the input regions we identified with retrograde tracing, the prelimbic area (PL), which has been shown to recruit cholinergic input to the cortex in rats^62–64^. We retrogradely expressed the red-shifted excitatory opsin ChRmine, first injected in the HDB, and placed optic fibers in the PL to stimulate the somata of HDB-projecting neurons (PL→HDB), ipsilateral to the injection. Note, however, that given that the light can potentially spread beyond PL, and that the underlying region (infralimbic area, ILA) also contains BF-projecting neurons, we adopt the more agnostic terminology mPFC→HDB.

First, we confirmed that mPFC→HDB neurons recruit cholinergic input to the cortex. We optogenetically stimulated the projection neurons while performing widefield imaging of cholinergic axons expressing the Ca^2+^ indicator jGCaMP8m, with isosbestic hemodynamic correction. We did this through a 5-mm cranial window implanted over the posterior dorsal cortex ipsilateral to the injection (**Fig. 6A, B**, n = 12 sessions from 2 ChAT-Cre x Flex-jGCaMP8m — note that this cortical territory predominantly receives HDB input and that this transgenic cross has negligible GCaMP expression in cortical ChAT+ interneurons^27^). While mice ran spontaneously on a spherical treadmill in a brightly lit environment, we used green light to stimulate mPFC→HDB neurons for 1 second (530-nm LED, 10 Hz, 1–1.2 mW, 10–20 trials with a random 90–120 s inter-trial interval). We determined that the brightly lit environment and the green excitation wavelength (as opposed to red) were necessary to minimize direct visual responses to the LED, in preliminary control experiments on animals not expressing the opsin (not shown). Thus, because 530 nm is also the emission wavelength of jGCaMP8m, we masked the imaging data during stimulation. Finally, we also used videos of one of the mouse’s eye and markerless tracking methods (Lightning Pose^65^) to measure pupil area during these experiments. We computed optogenetic-stimulation-triggered responses of cholinergic axons, averaged across the FOV. This revealed significant increases in cholinergic input to the cortex following mPFC→HDB stimulation (**Fig. 6C**, p = 4.88 × 10^-4^, signed rank test). This is compatible with the known monosynaptic targeting of BF cholinergic neurons by the PL^52^, though cholinergic activation could also be partly mediated via polysynaptic interactions within the BF^66,67^.

**Fig. 6.**
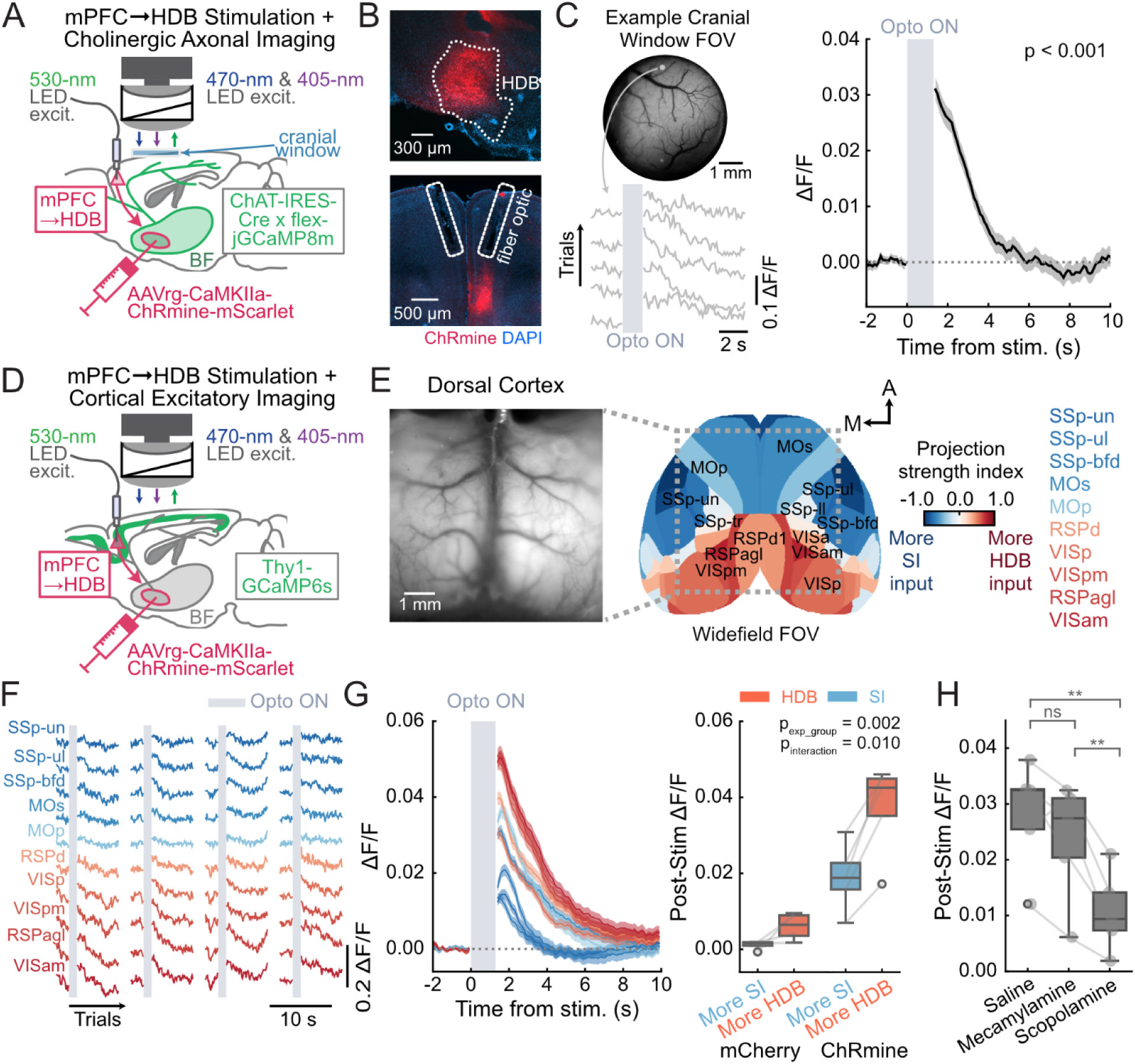
mPFC→HDB stimulation modulates cortical activity via a cholinergic mechanism. (A) Schematic of widefield Ca^2+^ imaging integrated with 530-nm LED stimulation for optogenetically activating the mPFC→HDB projection while imaging cholinergic axons in ChAT-IRES-Cre x flex-jGCaMP8m mice. (B) Left, fluorescence images of an example injection site in the HDB (outline hand-drawn from atlas) and optic fibers in PL. Note that, although we implanted fibers bilaterally, we only injected the virus unilaterally. (C) Top left, example cranial-window FOV for cholinergic-axon imaging. Bottom left, example single-trial ΔF/F traces aligned to mPFC→HDB stimulation, for the pixel indicated on top (arrow). Right, session average of mPFC→HDB stimulation-triggered activity of cholinergic axons, average across the whole FOV (n = 12 sessions from 2 mice). Shaded areas: ± S.E.M. across sessions. The printed p-value is from a Wilcoxon signed-rank test. (D) Same as A, for mPFC→HDB stimulation while imaging cortical excitatory neurons in Thy1-GCaMP6s mice. (E) Left, example cleared skull FOV from a Thy1-GCaMP6s mouse. Right, Allen CCFv3 parcellation of cortical regions within the FOV. Each region is color coded according to its relative strength of innervation by SI or HDB, given by the difference in projection density between HDB and SI, divided by its sum. (F) Example of single-trial stimulation-aligned ΔF/F, for pixels selected from each cortical region. (G) Left, average ΔF/F aligned to mPFC→HDB stimulation (n = 5 mice). Shaded areas: ± S.E.M. across trials (n = 365). The color scheme reflects the relative cholinergic projection strength from HDB (red) and SI (blue) as in panel E. Right, comparison of response magnitude (averaged between 0 and 1 s after stimulation offset) between dorsal cortical regions that receive predominantly HDB vs SI input. The printed p-values are from a two-way mixed ANOVA with factors experimental group (ChRmine, n = 5 mice vs. mCherry, n = 4 mice) and predominant input (average over HDB-receiving vs. SI receiving; we only selected areas with absolute projection strength index > 0.3). Lines: individual mice. (H) Stimulation-evoked responses averaged across dorsal cortical regions (computed during the same time window as in G) under vehicle or cholinergic-blocker administration (n = 5 mice). *: p < 0.05, **: p < 0.01, from post hoc paired t-tests. The tests followed a one-way repeated-measures ANOVA with drug as the factor (p = 8.27 × 10^-4^). Lines: individual mice.

Next, we sought to understand how this cholinergic recruitment potentially results in heterogeneous modulation of large-scale cortical activity. We performed similar experiments in mice expressing GCaMP6s in cortical excitatory neurons, using a cleared skull preparation to give us optical access to most of the dorsal cortex (**Fig. 6D**, n = 20 sessions from 5 Thy1-GCaMP6s mice). Our large FOV provided an opportunity to observe stimulation-induced changes in excitatory activity as a function of the relative predominance of HDB vs. SI input, which we quantified as a projection strength index using the anterograde data analyzed above (**Fig. 6E**). Activation of mPFC→HDB led to a cortex-wide increase in activity, roughly following the gradient of relative HDB innervation strength (**Fig. 6F, G**). Indeed, we found that, although all regions responded significantly (p < 0.001, Wilcoxon signed-rank test with FDR correction), predominantly HDB-receiving regions did so with larger magnitude (**Fig. 6G**, p = 3.44 × 10^-4^ for predominant input, two-way mixed ANOVA with factors experimental group and predominant input). This was not true for control animals not expressing the opsin (n = 4 Thy1-GCaMP6s mice injected with AAVrg-mCherry, p = 0.12, post-hoc t test with FDR correction). Further, experimental animals had significantly larger responses than controls (**Fig. 6G,** p = 0.01, interaction between experimental group and predominant input, two-way mixed ANOVA). This increase in cortical activity is compatible with previous reports of ACh-induced firing-rate increases in cortical neurons^24,68–70^. Moreover, although different running speeds and pupil-area dynamics were associated with slightly (but not significantly) different magnitudes of cortical responses to stimulation, the increase was overall robust to the animals’ behavioral state (**Fig. S7**). Finally, to probe whether such increases in cortical activity were mediated by cholinergic mechanisms (as suggested by **Fig. 6C**), we repeated these experiments with systemic (intraperitoneal) administration of scopolamine (0.5 mg/kg) and mecamylamine (1 mg/kg), a pan-muscarinic and a nicotinic antagonist, respectively (or vehicle saline as a control, n = 14 scopolamine and n = 17 mecamylamine sessions from the same 5 mice). We found that the magnitude of the effect of mPFC→HDB stimulation was significantly decreased by scopolamine (p = 4.63 × 10^-3^ vs. vehicle, significant effect in one-way repeated-measures ANOVA followed by post-hoc paired t-test), but not mecamylamine (p = 0.18 vs. vehicle), suggesting a predominantly muscarinic mechanism (**Fig. 6H**). However, scopolamine did not completely abolish the effects, which may have been due to incomplete antagonism or the additional recruitment of glutamatergic and/or GABAergic projections from the BF to the cortex. Overall, these results confirmed that activation of mPFC→HDB neurons recruits HDB projections to the cortex and modulates cortical activity via an at-least partly cholinergic mechanism.

### mPFC→HDB and mPFC→SI stimulation results in different spatial patterns of cortex-wide modulation

Finally, we asked whether stimulating mPFC→HDB or mPFC→SI produced distinct patterns of large-scale cortical modulation — which is not a given, since many mPFC neurons project to both subnuclei (**Fig. 5**). We performed similar experiments to the above in separate mice, targeting mPFC→SI neurons and imaging cortical excitatory activity (n = 9 sessions from 2 Thy1-GCaMP6s mice). On average, mPFC→HDB and mPFC→SI stimulation produced different spatial patterns, with peak activity at significantly more lateral cortical locations for mPFC→SI (**Fig. 7A, B**, p = 1.08 × 10^-6^, rank-sum test). This roughly matched the relative strength of HDB and SI input to the dorsal cortex (**Figs. 1C, 6E**). However, these analyses considered only the cortical location with peak activity, which was highly variable across trials (**Fig. 7B**). Thus, we wondered if more intricate spatial patterns of stimulation-induced cortical activity were also distinct between the stimulated projections. We trained L2-regularized logistic-regression models to classify (decode) the stimulated projection from the resulting pixel-wise activity patterns in the dorsal cortex in each trial (averaged at 0–1 s post-stimulation), first at a resolution of 150 µm/pixel (**Fig. 7C**). The decoders had notably high cross-validated accuracy (**Fig. 7D**, ∼95% overall), confirming that stimulating each projection led to very distinct spatial patterns of cortex-wide activity. To have more insight into what drove this distinction, we plotted the decoder weights on cortical space, which revealed patterns with sub-region resolution (**Fig. 7E**). Finally, to quantify the spatial scale at which activity induced by mPFC→HDB or mPFC→SI stimulation is maximally different, we spatially binned the cortical images at multiple resolutions (100–2000 µm/pixel) and again trained decoders for each. Interestingly, rather than smoothly decaying, decoding accuracy remained stably high up to a pixel size of 300–500 µm (i.e., smaller than the average cortical area), even when we subsampled pixels to match the number of model parameters across decoders (**Fig. 7F**). On the other hand, decoding accuracy was higher than chance even at the coarsest level, suggesting that the differences in cortical activity resulting from mPFC→HDB and mPFC→SI stimulation are robust and exist at multiple spatial scales.

**Fig. 7.**
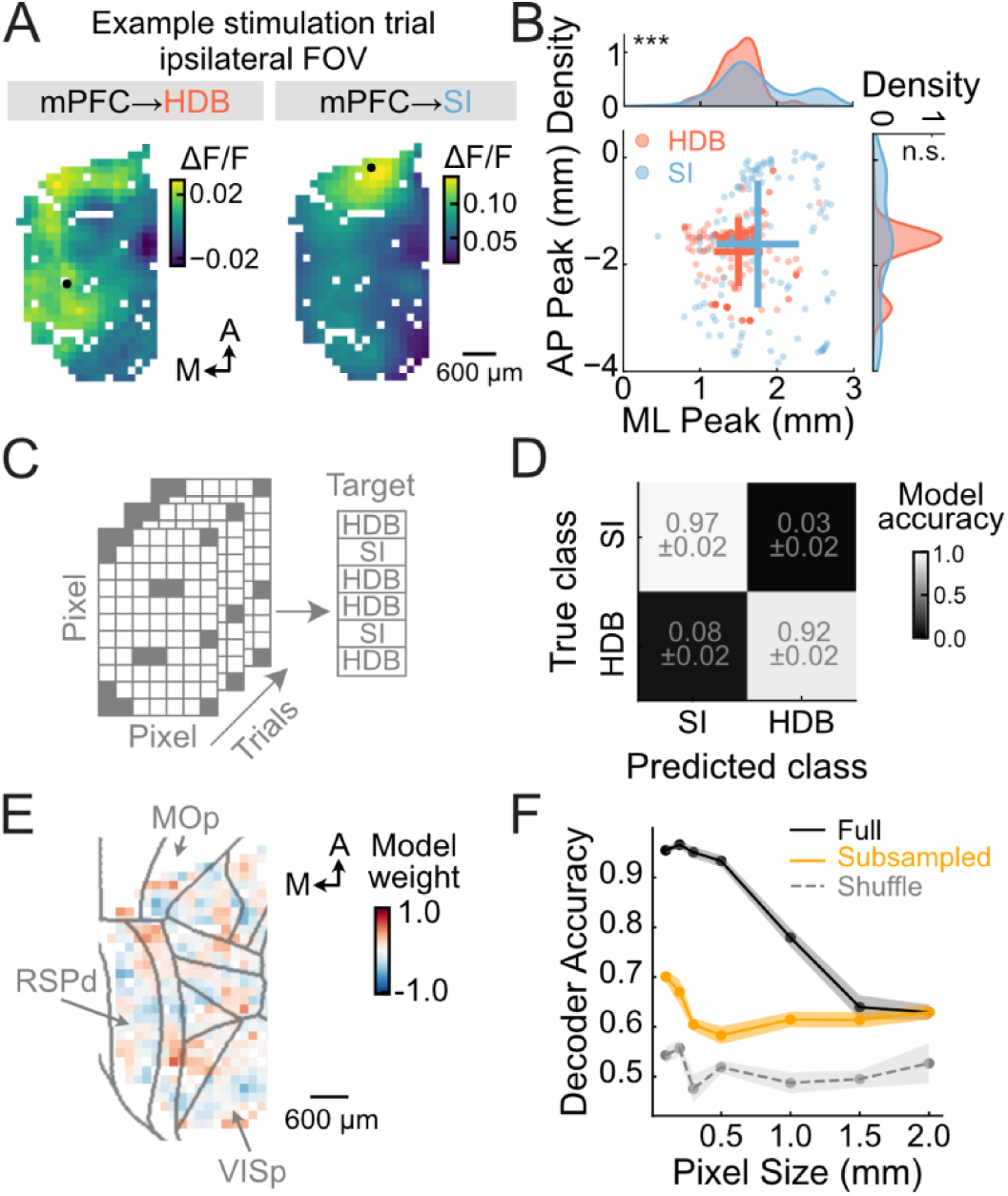
mPFC→HDB or mPFC→SI stimulation yields different cortex-wide activity patterns. (A) Example single-trial spatial maps of cortical excitatory ΔF/F, averaged over 1 s following the offset of optogenetic stimulation of mPFC→HDB (left) and mPFC→SI (right). Black dots on maps indicate the location of peak activity. (B) Distributions of peak-activity locations (AP and ML coordinates) across trials, for mPFC→HDB and mPFC→SI stimulation. Plots on top and to the right show the density of peaks along each anatomical axis. Crosses: mean ± S.D over trials. ***: p < 0.001, rank-sum test. (C) Schematic of the decoder trained to predict the stimulated projection from the pixel-wise cortical activity maps like the ones in panel A. The data were co-registered to the Allen CCFv3 so as to train a single model from different mice. (D) Confusion matrix showing decoder performance for a pixel size of 150 µm. (E) Average decoding weights over 5 cross-validation folds plotted on cortical space for the models in D. Gray lines indicate cortical-area borders. (F) Decoding accuracy as a function of pixel size. The ‘subsampled’ curve was obtained by randomly subsampling all decoders to match the one with the fewest parameters (2-mm pixels), so as to estimate the floor of performance for a given pixel size.

## Discussion

Using anterograde and retrograde tracing, we have shown that different mouse BF subnuclei preferentially target largely non-overlapping brain regions (**Figs. 1, 2**) and receive only partially overlapping input from a sparse set of brain regions (**Figs. 3–5**). Further, we combined cortex-wide widefield Ca^2+^ imaging with projection-specific optogenetic stimulation to show that this anatomical arrangement enables the independent recruitment of sub-area-specific patterns of cholinergic modulation of the dorsal cortex (**Figs. 6, 7**).

Our results are broadly compatible with previous anatomical characterization of the rodent BF^36,37,39,41,50,52–55^ (see **Table S2** for a comparison with previous mouse work), but they provide, to our knowledge, the first systematic comparison between mouse BF subnuclei using a consistent methodology, within the same study. Moreover, previous work specifically on the input-output relationship of the BF had primarily focused on characterizing which inputs cholinergic neurons projecting to specific cortical areas receive^36,37^. These studies generally found modular forebrain-BF-forebrain connectivity schemes, e.g. with BF cholinergic neurons projecting to the motor cortex receiving most input from the somatosensory cortex and basal ganglia^36^. Our results extend these findings by providing a broader, zoomed-out characterization at the level of BF subnuclei. They reveal a striking divergent organization whereby similar areas (though not necessarily neurons) project to BF subnuclei, which then broadcast to much larger territories with less overlap.

Our study also directly links this large-scale anatomical arrangement to spatially specific, large-scale cholinergic modulation of the dorsal cortex. Interestingly, both the overall pattern of anatomical segregation of HDB and SI input (**Fig. 6E**) and the effects on mPFC→HDB stimulation on cortical activity (**Fig. 6G**) closely mirror task-related differences in cortex-wide cholinergic-axon activity in mice switching between two decision-making tasks^27^. This suggests that the modulation patterns we measured in naïve mice running spontaneously could have relevance for understanding cholinergic modulation in the context of cognitive behaviors. It also suggests that task-specific recruitment of BF cholinergic neurons could in part come from mPFC projections to different BF subnuclei, which would be compatible with mPFC’s role in decision making and task switching in rodents^71–75^. Future studies should test this directly in the context of behavioral tasks. More broadly, future studies should also probe the behavioral role of partly segregated input from other forebrain regions (e.g., anterior insula and amygdala) to BF subnuclei.

Overall, our results outline an input-output anatomical arrangement that could underlie the behavior-specific recruitment of spatially heterogeneous cholinergic modulation^6,19,26,27,49^ that is thought to mediate ACh’s myriad functions^38,40^.

## Limitations of the study

Our retrograde viral strategy lacks cell-type specificity. First, this means we cannot be certain that the input patterns we observed (**Figs. 3–5**) can be directly mapped onto the cholinergic output of the BF subnuclei (**Figs. 1, 2**) — though note that different cell types in the BF do receive input from the same brain regions^52^. Second, because of this limitation, the mPFC→BF stimulation may also directly activate non-cholinergic BF neurons (cortically projecting or otherwise). In fact, a recent study found that parvalbumin-positive (PV) inhibitory neurons in the BF project to the cortex with a similar topology as cholinergic ones^76^. Because they preferentially target cortical PV interneurons, activating BF PV projections via mPFC→BF stimulation would have a disinhibitory effect on cortical excitatory neurons^61,76^. This could explain the residual effects of mPFC→BF stimulation we observed under cholinergic-receptor blockers (**Fig. 6H**). Moreover, it is possible that the effects of stimulating mPFC→BF projections on cortical activity via direct mPFC→dorsal-cortex activation, if mPFC→BF neurons collateralize. Future studies should characterize the collateralization patterns (if any) of these projection neurons. Finally, another limitation that should be addressed by future studies is our focus on only two of the major BF subnuclei, which also include the medial septum, the vertical limb of the diagonal band of Broca and the nucleus basalis magnocellularis.

## Data Accessibility

The full dataset will be publicly released upon publication of this work.

## Code Accessibility

The code will be publicly released upon publication of this work.

## Acknowledgements

This work was supported by the CAREER award IOS-2609476 from the National Science Foundation (L.P.), a McKnight Neuroscience Scholar Award (L.P.), and an NIH T32 to J.T. and R.L. (parent award 5T32MH067564). We thank Daniel Dombeck, Tiffany Schmidt, Yevgenia Kozorovitskiy, and Pinto lab members for excellent suggestions on figures and analyses. Ex-vivo microscopy was performed at the Northwestern University Center for Advanced Microscopy (RRID: SCR_020996) supported by NCI CCSG P30 CA060553 awarded to the Robert H Lurie Comprehensive Cancer Center, and at the University of Chicago Integrated Light Microscopy Core (RRID: SCR_019197) supported by the Cancer Center Support Grant (2P30CA014599).

## Author contributions

J.T. and L.P. conceived the project. J.T. and R.L. analyzed the anterograde tracing dataset. J.T. and M.K. performed the retrograde tracing experiments. J.T. and A.M.H. performed the widefield imaging and optogenetic experiments. J.T. analyzed the retrograde tracing and widefield imaging data. J.T. and L.P. wrote the manuscript. L.P. supervised the work.

## STAR Methods

### Key Resources Table

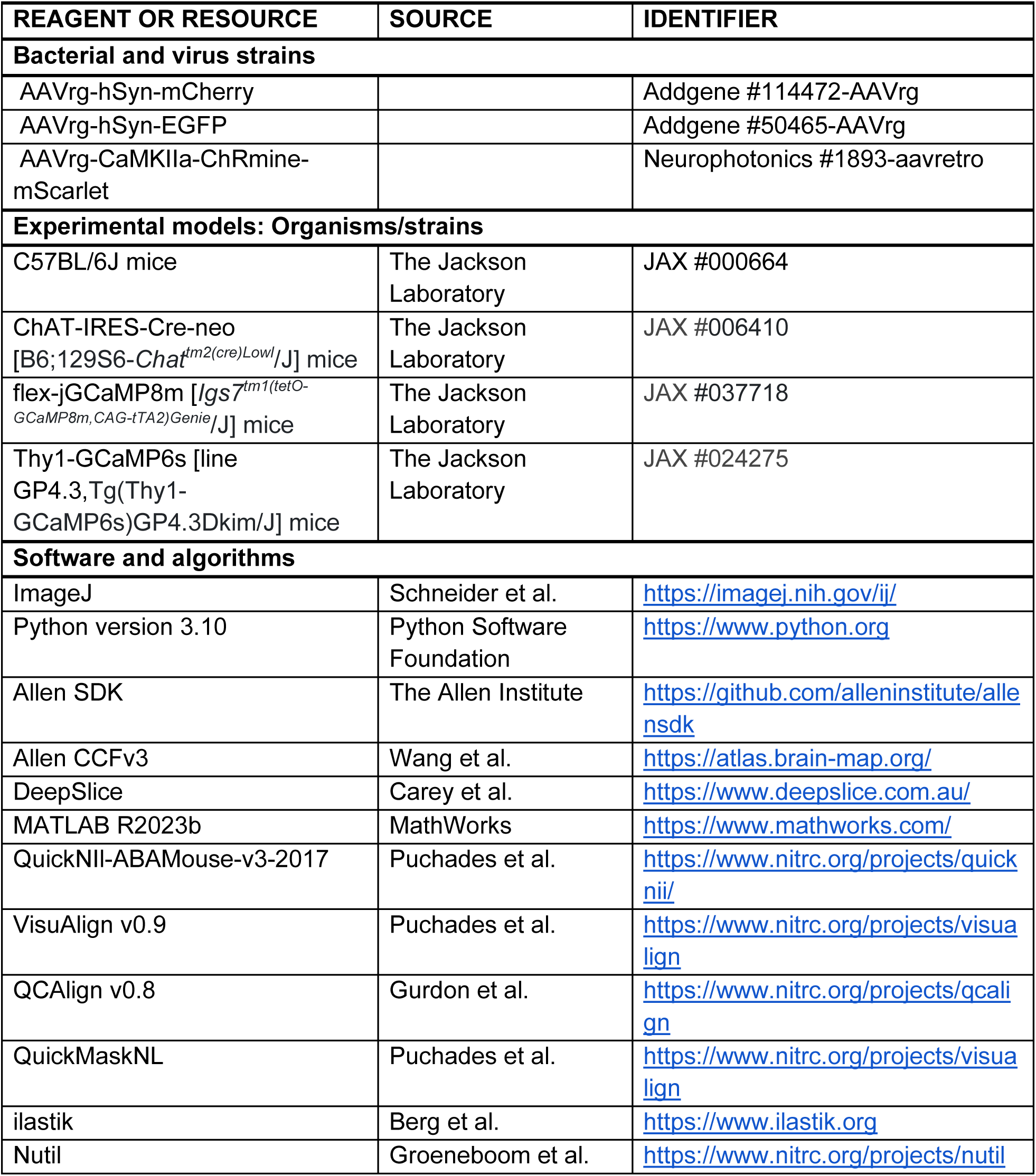

| **REAGENT OR RESOURCE** | **SOURCE** | **IDENTIFIER** |
|---|---|---|
| **Bacterial and virus strains** |  |  |
| AAVrg-hSyn-mCherry |  | Addgene #114472-AAVrg |
| AAVrg-hSyn-EGFP |  | Addgene #50465-AAVrg |
| AAVrg-CaMKIIa-ChRmine-mScarlet |  | Neurophotonics #1893-aavretro |
| **Experimental models: Organisms/strains** |  |  |
| C57BL/6J mice | The Jackson Laboratory | JAX #000664 |
| ChAT-IRES-Cre-neo [B6;129S6-*Chat*<sup>*tm2(cre)Lowl*</sup>/J] mice | The Jackson Laboratory | JAX #006410 |
| flex-jGCaMP8m [*Igs7*<sup>*tm1(tetO-GCaMP8m,CAG-tTA2)Genie*</sup>/J] mice | The Jackson Laboratory | JAX #037718 |
| Thy1-GCaMP6s [line GP4.3,Tg(Thy1-GCaMP6s)GP4.3Dkim/J] mice | The Jackson Laboratory | JAX #024275 |
| **Software and algorithms** |  |  |
| ImageJ | Schneider et al. | https://imagej.nih.gov/ij/ |
| Python version 3.10 | Python Software Foundation | https://www.python.org |
| Allen SDK | The Allen Institute | https://github.com/alleninstitute/allensdk |
| Allen CCFv3 | Wang et al. | https://atlas.brain-map.org/ |
| DeepSlice | Carey et al. | https://www.deepslice.com.au/ |
| MATLAB R2023b | MathWorks | https://www.mathworks.com/ |
| QuickNII-ABAMouse-v3-2017 | Puchades et al. | https://www.nitrc.org/projects/quicknii/ |
| VisuAlign v0.9 | Puchades et al. | https://www.nitrc.org/projects/visualign |
| QCAlign v0.8 | Gurdon et al. | https://www.nitrc.org/projects/qcalign |
| QuickMaskNL | Puchades et al. | https://www.nitrc.org/projects/visualign |
| ilastik | Berg et al. | https://www.ilastik.org |
| Nutil | Groeneboom et al. | https://www.nitrc.org/projects/nutil |

### Animals

All experimental procedures were approved by Northwestern University’s and The University of Chicago’s Institutional Animal Care and Use Committees, and were done in accordance with the Guide for the Care and Use of Laboratory Animals. Mice were kept under a 12/12-h reverse light/dark cycle with ad libitum access to food and water. We used a total of 21 mice of various genotypes and both sexes, aged 2–14 months at the time of data collection, as detailed below.

● Retrograde viral tracing in **Figs. 3–5, S5, S6**. A total of 8 C57BL/6J mice [JAX stock #000664] were used: 4 for injection in the HDB; 6 for injection in the SI (3 of these have injections for both); 1 for dual-color injections in HDB to estimate the efficiency ceiling.
● Cholinergic-axon imaging experiments in **Fig. 6**. Crosses between ChAT-IRES-Cre-neo [B6;129S6-*Chat^tm2(cre)Lowl^*/J, JAX stock #006410] and flex-jGCaMP8m [*Igs7^tm1(tetO-GCaMP8m,CAG-tTA2)Genie^*/J, JAX stock #037718] (n = 2).
● Cortical excitatory neuron imaging experiments in **Figs. 6, 7, S7**. Thy1-GCaMP6s [line GP4.3,Tg(Thy1-GCaMP6s)GP4.3Dkim/J, JAX stock # 024275] (n = 11).
  ○ Mice with retrograde ChRmine injected in the HDB (n = 5).
  ○ Mice with retrograde ChRmine injected in the SI (n = 2).
  ○ Mice with retrograde mCherry injected in the HDB (n = 4).

For widefield imaging experiments (**Figs. 6, 7**), mice recovered for at least 5 days after surgery before undergoing habituation to handling and the running apparatus. Following each imaging session, the mice were given access to an enclosure containing toys and conspecifics for environmental enrichment.

### Surgical procedures

Mice aged 8–12 weeks underwent a single surgical procedure that varied depending on the experiment, as described below. In all procedures, mice were first anesthetized using isoflurane in oxygen (3% v/v induction, 1.5% maintenance) and were secured in a stereotaxic frame (Kopf). After trichotomy and asepsis, we performed an incision on the scalp to expose the dorsal skull.

For dual retrograde tracing experiments (**Figs. 3, 4**), we performed intracranial injections to deliver viral retrograde tracers (retrograde AAV, AAVrg) in the two BF subnuclei, HDB and SI, within the same animal or separately (EGFP or mCherry, counterbalanced across subnuclei). We made one or two small craniotomies (∼500-μm diameter) using an air-driven dental drill with a carbide burr to target the HDB (AP: 0.6 mm from bregma, ML: 2.17 mm, DV: -5 mm) and/or the SI (AP: -0.23 mm, ML: 2.6 mm, DV: -4.48 mm), both in the right hemisphere and with the microsyringe inserted at a 15-degree angle. We delivered 80–100 nL of either AAVrg-hSyn-mCherry (Addgene #114472-AAVrg, titer 7 × 10^12^) or AAVrg-hSyn-EGFP (Addgene #50465-AAVrg, titer 7 × 10^12^) into each subnuclei using a microsyringe pump at a rate of 75 nL/min (World Precision Instruments). We waited for 10–15 minutes before retracting the syringe. These parameters allowed us to target a ∼1-mm-diameter region around the injection site, avoiding overlap between HDB and SI, as determined with preliminary experiments (not shown). To estimate the ceiling of viral expression efficiency, we also injected one animal with a mixture of both tracers injected in the same subnucleus. After the injection, the craniotomy was sealed with bone wax (Surgical Specialties), and the scalp was closed using 5-0 Nylon sutures (Med Vet). Lastly, topical triple antibiotic ointment was applied on the scalp. Following 3–4 weeks of viral incubation, the mice were perfused and the brains were extracted for histology as described below.

For simultaneous widefield imaging and optogenetic stimulation experiments (**Figs. 6, 7**), a small craniotomy was first made to target either of the HDB or SI in the right hemisphere (coordinates as above, only one subnucleus per animal). We then virally delivered the excitatory opsin ChRmine with an intracranial injection as above (AAVrg-CaMKIIa-ChRmine-mScarlet, Neurophotonics #1893-aavretro, titer 9.7 × 10^12^, 100 nL, 50–75 nL/min. The craniotomy was then sealed with bone wax (Surgical Specialties). After the injection, we implanted fiber optics to target HDB-or SI-projecting neurons in the prelimbic cortex (PL). Two more small craniotomies were made and the fibers were implanted bilaterally at a 15-degree angle from the vertical plane, with the fiber tip sitting 0.5 mm above the dorsal border of PL (AP: 2 mm, ML: 0.9 mm, DV -1.3 mm). The fiber-ferrule assemblies were secured using a thin layer of cyanoacrylate adhesive (3M Vetbond) to seal the craniotomy and another layer of clear metabond (Parkell) to secure the ferrule base with the surrounding skull. We left ∼0.7-mm length of ferrule end exposed outside for optic fiber patch connection. Once the cement around both fiber assemblies was cured, we began the cranial window (**Fig. 6A**) or cleared skull (**Fig. 6C**) procedures. For the cranial window, using a dental drill and fine forceps, we first created a 5-mm diameter circular craniotomy in the right hemisphere centered around AP -2 mm and ML -2 mm. We then lowered the circular glass window onto the craniotomy using a manipulator and bonded it to the skull using veterinary cyanoacrylate glue and metabond (Parkell). For the cleared skull preparation, we followed procedures based on previously described protocols^27,77^. The periosteum over the dorsal skull was gently scraped off before applying a thin layer of cyanoacrylate adhesive and clear metabond on top. After curing, the metabond surface was smoothed using a cement polishing kit (Pearson Dental), and a titanium headplate implant was affixed to the skull with opaque metabond (Parkell) and painted over with a thin layer of carbon powder and opaque metabond mix to block off light from the fiber optic. Lastly, a uniform, thin coat of transparent nail polish (Electron Microscope Science) was applied across the implant.

For analgesia, mice received a buprenorphine ER (1 mg/kg S.C.) or Ethiqa XR (3.25 mg/kg S.C.) before the surgery, as well as meloxicam (20 mg/kg S.C.) before and 24 hours after the surgery. They also received warm saline (0.1–0.5 mL S.C.) during and after the procedure to maintain hydration.

### Transcardial perfusion, histology and *ex-vivo* imaging

Mice were deeply anesthetized with ketamine (50–100 mg/kg) and xylazine (10 mg/kg) and transcardially perfused with phosphate-buffered saline (PBS, 20 mL), followed by 4% paraformaldehyde (10 mL). The brains were then extracted and post-fixed in 4% w/v paraformaldehyde for 12–24 h, and then cryoprotected with 30% w/v sucrose. The tissue was sectioned coronally in 50-µm slices with a cryostat (Leica), and every third section through the forebrain was mounted on Superfrost slides (Thermo Fisher) with Fluoromount-G^TM^ mounting medium with DAPI (Invitrogen) and then coverslipped.

To quantify the expression of retrograde tracers, widefield images were obtained on a Nikon Ti2 Widefield (4× magnification) and/or Zeiss Axio Imager 2 (5×). Images were stitched using Nikon software (NIS Element) to produce large images of the coronal slice. These large images were then registered to atlas and quantified using procedures described below.

To quantify the same-neuron overlap of input to the two subnuclei coming from a few selected regions (**Fig. 5**), we obtained confocal z-stacks on a Leica SP8 confocal microscope using a 40× magnification oil immersion objective. Cell counts were performed manually using Fiji by two experimenters and averaged.

### Mesoscale widefield Ca^2+^ imaging and eye tracking

We used a custom-built apparatus built for head-fixed mice running on an air-suspended Styrofoam© and surrounded by a spherical screen, described in detail elsewhere^78^. The mice ran freely under uniform white light projected onto the screen, and running velocity was recorded using an optical velocity sensor underneath the ball, connected to an Arduino Due. The behavioral apparatus was integrated with a custom-built widefield macroscope, consisting of 1×–0.63× tandem lenses (Leica) with a dichroic beamsplitter and green emission filter (Semrock, FF495-DI03-50X70 and FF01-525/45), a Prime BSI Express sCMOS (Teledyne), and alternating illumination with blue and violet LEDs (Cairn, 470 nm and 405 nm respectively, 40-nm bandpass filtered, ∼2 mW/cm^2^ power at the sample). Images were captured at 20 Hz with 1024 × 1024 pixel resolution (6.8-µm pixel size, ∼ 7 × 7-mm field of view), controlled by µManager on a PC. We synchronized imaging with behavioral data by routing sCMOS exposure signals through a National Instruments DAQ card to the behavior PC, saving the signals directly to the behavioral data file. We also performed eye tracking by filming a mirror positioned at an angle under the mouse’s left eye. We used a CMOS blackfly camera (Teledyne, 1920 × 1200 pixels) with a 35-mm focal-length lens (Tamron M112FM) and an infra-red LED with a broadband diffuser (740 nm, Digikey) behind the mirror, driven by a custom-printed circuit board to help illuminate the recording. Images were acquired at 15 Hz using a second instance of μManager running on the same PC as the widefield camera, and synchronized by saving exposure signals into the behavioral data file as above.

### Optogenetics and systemic pharmacology

For optogenetic experiments, optic fiber assemblies (Thorlab 200-μm core, 0.39 NA, FT200UMT) were manually fabricated and cut 1.5 mm below the ferrule tip (Precision Fiber Products LC 1.25-mm OD Multimode Ceramic Zirconia Ferrules, MM-FER2007C, 1.4-mm length). Fiber placement was verified histologically for all animals. For stimulation, 530-nm excitation LED light (Prizmatic) was delivered bilaterally to the PL through a 200-µm optical fiber patch cable, with illumination power of 1–1.2 mW at the fiber tip (∼0.7–0.85 mW/mm^2^ measured ∼0.5 mm below the fiber tip; though notice only one hemisphere received a viral injection). The LED was pulsed in a square wave at 10 Hz for 1 s at 50% duty cycle for 10–20 trials. Intertrial intervals were varied in length from 90–120 s, drawn from a uniform distribution. The LED was controlled by the VR software using TTL pulses. Given the proximity of the mPFC to the retina, to minimize the visual response to the LED pulses, the retina was adapted with bright white light projected onto the screen. This was determined in preliminary experiments based on the parameters that minimized visual cortical responses in opsin-negative mice.

For experiments involving pharmacology, the mice were injected with either scopolamine (0.5 mg/kg in saline), mecamylamine (1 mg/kg in saline) or vehicle (saline) intraperitoneally, 30 min before the start of the behavioral session. Each animal underwent 3 sessions for each of the 3 treatments, with order determined randomly.

### Analysis of anterograde tracing data from the Allen Mouse Connectivity Atlas

Anterograde tracing data (**Figs. 1, 2, S1–S4**) were obtained from the Allen Institute Mouse Connectivity Atlas using custom-written Python software to interface with the Allen SDK. We first selected injections with EGFP or SynEGFP fluorophore in the HDB and SI of ChAT-IRES-Cre-neo mice. Because HDB and SI are both curved, elongated structures along the AP axis, to ensure consistency and lack of overlap between subnuclei, we selected a subset of injections that were ± 0.5 mm around AP 0.6 mm for HDB and AP -0.3 mm for SI, similar to the target coordinates we used for the injection of retrograde tracers (though here we did not impose restrictions on ML and DV coordinates). This resulted in 3 mice for HDB and 4 for SI, corresponding to Allen experiment IDs [167200755, 177605425, 504173156] for HDB and [478491810, 126841788, 478581786, 504521113] for SI.

We used the Allen-provided projection signal, determined using a signal-detection algorithm to segment fluorescent signals from the background. Voxel-level projection density was defined per 25-µm × 25-µm × 25-µm voxel and computed as the sum of detected projection pixels divided by sum of all pixels in voxel^56^. All voxels were first filtered using the provided binary mask to remove off-brain fluorescence for each experiment. For all subsequent analyses, we additionally excluded structures at or surrounding the injection site: HDB, SI, medial septal nucleus, magnocellular nucleus, hypothalamic lateral zone, medial and median preoptic nuclei.

For voxel-level analyses (**Figs. 1, 2A, S1, S2, Table S1**), projection densities across voxels were normalized to the maximum projection density within each experiment, followed by averaging across experiments (**Fig. 1B**). To compare the projection between the two subnuclei along different anatomical planes (horizontal, coronal, sagittal), we maximum-projected the experiment-average projection densities along each plane for each subnucleus (**Fig. S1**). We then plotted the difference between these max-projected projection densities from injections in HDB and SI (**Fig. 1C**). To further quantify these differences along the planes, we averaged projection density over one axis at a time and plotted the experiment-averaged projection densities along the anteroposterior (AP), mediolateral (ML) and dorsoventral (DV) anatomical axes (**Figs. 1D, S2**). To generate visualizations of projection density across coronal volumes for each subnucleus (**Fig. 2A**), we first segmented projection volume in 1-mm coronal volumes between 2 and -3 mm AP (relative to bregma), then averaged the projection density across the AP axis. To test for whether the distributions of SI and HDB densities along each anatomical axis were different, we fitted a full model using ordinary least squares (OLS) regression to predict z-scored projection density *D* as a function of subnucleus identity, *id*, and a 3rd-degree polynomial expansion of the axis coordinates *x*, as well as the interactions between them:

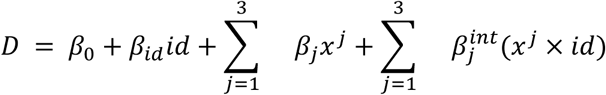

where *id* is a one-hot vector with 1’s for HDB and 0’s otherwise, *j* is the degree of the polynomial, the superscript *int* indicates the interaction term, and *x* ∈ {*ml*, *ap*, *dv*}. The models provided good approximations of the data (for ML, *R*^2^ = 0.688; for AP, *R*^2^ = 0.685; for DV, *R*^2^ = 0.587). We then fitted a nested reduced model excluding all the interaction terms and compared the performance of the full and reduced models using an F-test, with a significant value indicating that the interactions capture a significant amount of variance.

For brain-region-level analyses (**Fig. 2B, Table S1**), we averaged normalized voxel-level projection densities for each region, combining any layer and subnucleus of the structures. We then averaged these regional projection densities across experiments for each injection site. To test for significance (**Fig. 2B** only), we jointly modeled z-scored projection density for all regions and both subnuclei using OLS regression with subnucleus identity *id*, forebrain regions *r_j_* (each a categorical variable), and their interaction *int* as predictors:

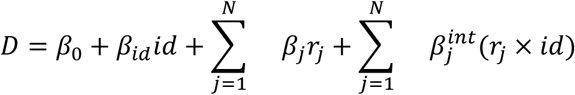

where *N* is the number of cerebrum regions (89). The model had good performance (*R*^2^ = 0.566). We first ran a Wald test to identify whether the region terms *r_j_*or the interaction terms were collectively significant. Then, for each region, we computed the coefficient contrast as the model-predicted difference between injection locations, and used a *t* distribution to compute p-values, with false discovery rate (FDR) correction.

For the analysis in **Fig. S3**, we segmented the data in 0.5-mm coronal volumes between 3 to -4.5 mm AP relative to bregma. Using the reference annotation of voxel labels provided by the Allen SDK, we obtained a list of brain regions within each volume. Then, for each brain region in the volume, we averaged the projection density within its voxels inside the volume. Finally, for analysis of cortical layers (**Fig. S4**), we excluded layer 4, since it is not present in all cortical areas. We then computed the average layer projection density across cortical regions and experiments (**Fig. S4B**), as well as the average proportion of projection density for each layer across individual regions (**Fig. S4C**).

### Brain atlas alignment and automated detection of retrograde tracers

For most retrograde tracing experiments (**Figs. 3, 4, S5, S6, Table S1**), epifluorescence images of coronal brain slices were aligned to the Allen Mouse CCFv3^58^ following the QUINT workflow^60^. Images were first manually preprocessed in Fiji, adjusting brightness to cap the maximum value of the look-up table (LUT) histogram at 12000 for green and 18000 for red channels, respectively, then rotating and cropping the images. Images were downsampled to 40% of their native pixel resolution using Nutil Resize (final pixel size of 2.65 µm) for faster subsequent pixel classification^79^. We used the DAPI channel for atlas registration. These images were sorted by their AP coordinate and uploaded to DeepSlice^80^, an automatic mouse brain slice registration to the Allen CCFv3, with QuickNII used for manual fine adjustment to match the reference atlas^81^. We then exported the results to VisuAlign for in-plane nonlinear registration to the atlas, stretching reference atlas borders and moving landmark regions to match their locations on the brain slices^81,82^. Next, we used QuickMaskNL to generate hemibrain masks on atlas alignment to exclude the contralateral hemisphere^81,82^. We used an artificial-intelligence-generated (Anthropic Claude Opus 5.5) Fiji macro to add injection sites to the hemibrain masks. For downstream analyses, we excluded the same set of injection structures used in anterograde dataset above. Any tissue damage on the slices was annotated with QCAlign^81,82^ and slices with more than 30% damage were removed from the dataset.

Next, we used illastik, a supervised pixel classification software, to detect the locations of retrogradely labeled cells, agnostically referred to as ‘puncta’^83^. We manually labeled training data on a subset of brain slices and trained the model to identify pixels of puncta as targets vs. non-target background pixels. We chose 3–5 coronal sections per brain at different AP positions as training images to sample the range of background variation and tracer intensity across the brain volume. For each image, we manually labeled a subset of puncta and background pixels by drawing labels on representative pixels of each class. A random forest classifier was trained in Ilastik using color intensity, edge, and texture features across all available scales (σ = 0.3–10 pixels). We manually inspected the predictions from the classifier in an interactive live mode and iteratively refined and added annotations of uncertain or incorrect classifications until predictions were visually consistent across all training images. Models for green and red fluorescence were trained separately. Fluorescence classification results were exported as binary images, where each pixel indicated whether it is part of a punctum and pixel locations were anchored to atlas locations with results from VisuAlign or QCAlign. Finally, we used another AI-generated Fiji macro to detect and split large puncta with more than 200 pixels into tiles with the average punctum size for that slice (16–20 pixels). Only puncta containing at least 10 pixels were selected for subsequent quantification. Puncta were counted using the Nutil Quantifier^79^.

### Analysis of retrograde-tracing data

Retrograde tracing data were checked for viral expression and location. We confirmed the injection sites for each experiment by finding the center of mass of the injection data (**Figs. 3B, S5**). We included experiments with centers located ± 0.5 mm around AP 0.6 mm, ML 0–2 mm, and DV 5–7 mm for HDB and AP -0.3 mm, ML 1–3 mm, and DV 4–6 mm for SI. Note that the AP ranges matched the selection criteria for anterograde tracing data. A total of 7 animals passed our criteria: 4 with HDB injections (2 EGFP and 2 mCherry) and 6 with SI injections (2 EGFP and 4 mCherry). Three of the 7 animals had both HDB and SI injections that were also used for confocal overlap quantification in **Fig. 5**.

For voxel-level analyses (**Figs. 3, 4A, S5**), we first mapped the 2-D spatial coordinates of the retrograde puncta obtained from the Nutil Quantifier onto the Allen CCFv3 space by creating an empty 3-D matrix of the 25-µm CCFv3 dimensions and registered each coordinate pair to the nearest voxel. We defined input density as the puncta count within each 25-µm voxel divided by the largest single-voxel count within a given brain. To quantify the spatial distribution of input puncta between two nuclei along the major anatomical axes (**Fig. 3C**), we averaged input density over one axis at a time and plotted the experiment average of input densities along each axis. To test for significance, we used an F-test to compare full and reduced OLS models as described for voxel-level analyses in anterograde-tracing data. To generate 2-D visualizations of input density across in coronal volumes for each subnucleus (**Fig. 4A**), we first segmented 3-D density volume in 1-mm coronal volumes between 2 and -3 mm AP from bregma, then averaged the input density across the anteroposterior axis.

For brain-region-level analyses (**Figs. 4B, S6, Table S1**), we defined regional input density as the ratio between the puncta counts within a region and the total count for each experiment. We then averaged the input density across experiments for each region and tested statistical significance using the same OLS, Wald test, and post-hoc t-test approach as for the region-level analyses of antegrade-tracing data. For the analysis of cortical layers, we excluded layers 1 and 4, since layer 1 did not contain projection neurons and layer 4 is not present in all cortical areas. We computed the proportion of layer input density within individual regions for few selected regions averaged across experiments (**Fig. S6A**) and the average proportion of layer input density across regions (**Fig. S6B**).

### Widefield data preprocessing and analysis of stimulation-evoked responses

We have described the preprocessing procedures in detail elsewhere^27,78^. We used the same procedures for both cholinergic axonal imaging through a cranial window (**Fig. 6A–C**) and cortical excitatory imaging through the cleared skull (**Fig. 6D–H**). In brief, we performed motion correction using phase correlation and then spatially binned the imaging frames to 128 × 128 pixels (pixel size of ∼54 μm). We corrected Ca^2+^-dependent GCaMP fluorescence captured in the blue channel (470 nm) for hemodynamic contamination using the isosbestic, Ca^2+^-independent violet channel (405 nm) as follows. For each trial of optogenetic stimulation, we defined baseline fluorescence *F*_0_ for each alternating excitation wavelength by averaging 2 s of raw fluorescence before stimulation onset. We then computed *ΔF*/*F*(*t*) separately for each channel, *ΔF*/*F_b_*(*t*) and *ΔF*/*F_v_*(t), by computing [*F*(*t*) − *F*_0_]/*F*_0_ − 1, and computed a scaled version of 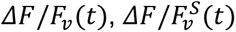, by finding the ratio between the median negative deflection of *ΔF*/*F_b_*(*t*) and *ΔF*/*F_v_*(*t*) and multiplying *ΔF*/*F_v_*(*t*) by this ratio. Because hemodynamic contamination is multiplicative^84^, we obtained *ΔF*/*F*(*t*) by dividing *ΔF*/*F_b_*(*t*) + 1 by 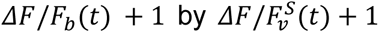 (for numerical stability) and then subtracting 1. For each session, we combined manual masking of the sagittal sinus and off-brain pixels (e.g. implant border, injection, and implant sites) with a vascular mask generated with adaptive median filtering^27^. Finally, we computed the signal-to-noise ratio (SNR) for each pixel using *ΔF*/*F*(*t*) from the 30 s preceding stimulation onset, concatenated across trials. We defined SNR as the ratio between a low-pass-filtered (i.e. signal) and a high-pass-filtered (i.e. noise) version of the *ΔF*/*F*(*t*) (3-Hz cutoff, order-10 Butterworth filter for both)^78^. We excluded pixels with SNR < 6 by incorporating them into the vascular mask.

For sessions imaging cholinergic axonal activity through cranial windows (**Fig. 6A–C**), we report the FOV-averaged *ΔF*/*F*(*t*) after applying the combined mask. For sessions imaging cortical excitatory activity across the dorsal cortex (**Fig. 6D–H**), we registered each session’s average imaging frame to the Allen CCFv3 space for downstream analyses^78^. We defined a reference image for each animal and manually marked the bregma and lambda on it. Then, for each session, we first used a Matlab image intensity-based optimizer (‘imregister’) and estimated the geometric transformation (‘imregtform’) to register its average image to the reference image. We then computed a second rigid transformation matrix to align bregma and lambda from the reference image to the bregma and lambda in CCF space.

For the cortical-region-level analyses in **Figs. 6** and **S7**, we assigned each pixel coordinate to a cortical region on the brain surface by detecting the first voxel with an area label along the DV axis in the reference annotation volume from the Allen SDK. Finally, using these pixel-wise area labels, we averaged *ΔF*/*F*(*t*) across pixels within each area and again averaged across trials from different sessions and animals for 10 s after stimulation onset (excluding 0–1 s given light contamination by the optogenetics LED). Standard errors in **Figs. 6C, G** and **S7A, C** were computed across all trials for display only. For statistical analysis of post-stimulus *ΔF*/*F*, we averaged *ΔF*/*F*(*t*) for the 1-s period immediately following the offset of stimulation, individually for each animal.

For the pixel-level analyses in **Fig. 7A, B**, we converted pixel coordinates into voxels in CCFv3 space by creating an empty 2-D matrix of the 25-um resolution CCFv3 space in the AP and ML dimensions and registering each coordinate pair to the nearest voxel location. For pixel-level post-stimulation *ΔF*/*F*, we applied a spatial binning factor of 6 (150-um) to avoid sparsity in the CCFv3 space and averaged *ΔF*/*F* from coordinate pairs assigned to each voxel (**Fig. 7A–C**). To identify the peak location for each trial, we smoothed the post-stimulus *ΔF*/*F* pixel map with a gaussian filter with a standard deviation of 0.5 pixels, and defined the peak location as the mean coordinate of the pixels above the 95^th^ percentile of the smoothed *ΔF*/*F* distribution. We then computed the kernel density estimate histograms with Gaussian smoothing with bandwidth determined by Scott’s Rule.

### Running analysis and pupilometry

For running analysis (**Fig. S7A, B**), the speed of the mouse was computed using the Euclidean norm of the x-and y-displacement of the ball over each widefield imaging frame (100 ms per frame after hemodynamic correction). Speed traces were smoothed with a moving average of 5 frames, z-scored using the mean and standard deviation of the whole-session traces, and aligned to optogenetic-stimulation trials. We then averaged the z-scored running speed 1 s before and 1 s after stimulation onset. We classified trials as ‘Runs’ if the average post-stimulation speed was greater than the baseline, and ‘Stops’ otherwise.

To track the mouse’s pupil dynamics (**Fig. S7C, D**), we used the markerless tracking algorithm Lightning Pose^65^ as we have described in detail elsewhere^27^. Briefly, after extracting the coordinates for 8 markers around the diameter of the pupil, for each frame of the eye movies, we fitted an ellipse to the markers and computed pupil area in squared pixels from the fitted major and minor axes. We excluded frames with a marker-average likelihood below 0.7 (estimated by Lighting Pose), since they typically correspond to occlusion of the eyes by the mouse’s body or blinking. After exclusion, pupil area traces for each session were interpolated linearly to fill in the excluded frames, smoothed with a moving average of 5 frames, z-scored using the mean and standard deviation of the whole-session traces, and aligned to stimulation trials. Trials that had more than 50% of excluded frames (determined prior to interpolation) were excluded from the analysis. Finally, we averaged z-scored pupil area 1 s before and after the onset of stimulation. We classified trials as ‘Baseline Constricted’ if the average post-stimulation area was greater than the baseline, and ‘Baseline Dilated’ otherwise.

### Decoding model

For all mPFC→HDB and mPFC→SI stimulation trials, pixelwise post-stimulus *ΔF*/*F* was binned with a factor of 6 to a pixel size of 150 µm x 150 µm and aligned across sessions in the Allen CCF space. We computed a common mask by identifying and combining locations of NaN pixels (from masking) across trials in each stimulation type, so as to remove any invalid pixels across sessions. Valid pixels were then arranged into a matrix of dimensions pixels × trials as predictors, balanced by resampling to the number of trials in the smaller (SI) class, into a L2-regularized, logistic regression using the sklearn package in Python. The stimulated projections were passed in as targets. We used stratified 5-fold cross-validation, and assessed accuracy on held out data. Model weights were averaged across folds. For the analysis comparing model performance with varying pixel size (**Fig. 7F**), we used the same regression method for each pixel-binning level (100, 200, 300, 500, 1000, 1500, and 2000 µm). To control for the fact that models with larger pixel size contained fewer parameters, we performed the same regression for each pixel size by randomly undersampling predictors in larger models to match the smallest model (n = 3 pixels), once per cross-validation fold. Finally, to obtain shuffle decoder performance, we randomly shuffled the target labels across the entire dataset once per cross-validation fold.

### General statistics

We detail statistical tests using OLS regression in their respective sections. For pairwise tests, we first assessed normality with the Shapiro-Wilk test to select between parametric (t-test) or non-parametric (rank-sum) tests, or their paired equivalents. For comparisons between multiple groups, we applied an analysis of variance (ANOVA), one-or two-way, with or without repeated measures, or mixed) followed by a post-hoc paired t-test comparison within groups. Whenever multiple comparisons were performed, we applied FDR correction to adjust p-values.

## Supplementary Information

**Fig. S1.**
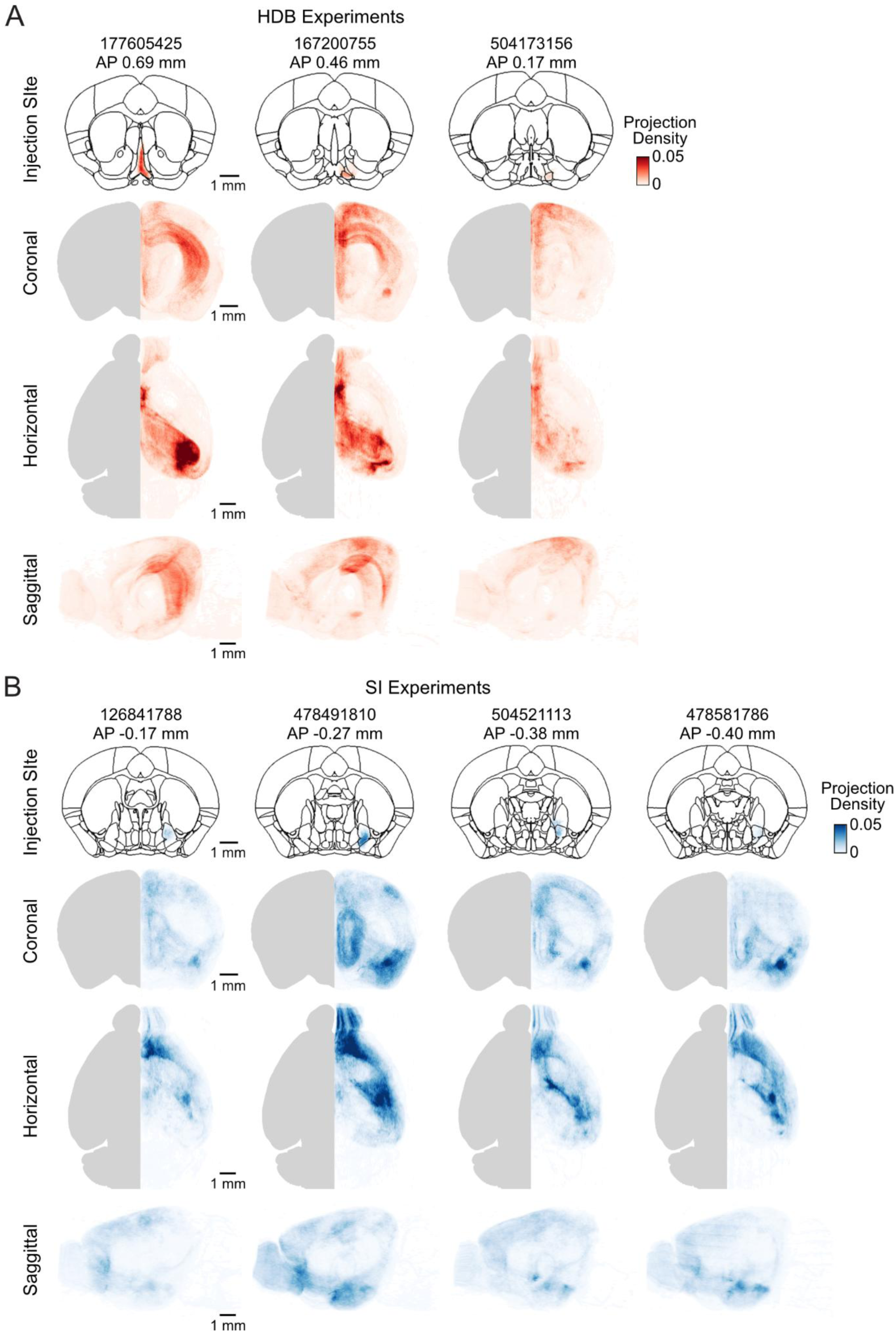
Injection site and cholinergic projections from individual experiments. (A) Top, injection locations for individual mice in the HDB dataset, with corresponding experiment IDs from the Allen Mouse Connectivity Atlas. Bottom, average cholinergic-projection density onto three anatomical planes for each brain. (B) Same as A, for the SI dataset.

**Fig. S2.**
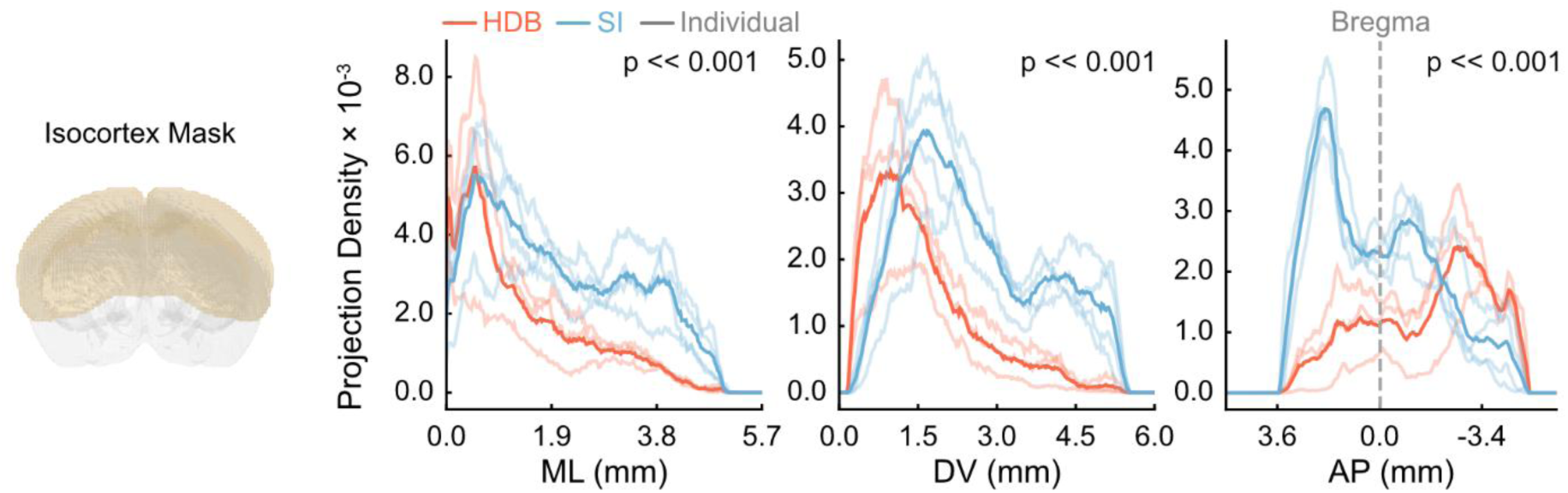
1-D spatial profiles of cholinergic projection density from each subnucleus across the isocortex, for individual mice and their averages. The printed p-values are from F-test for nested linear-regression models with and without interaction terms between injection location and projection density (see STAR methods). For ML, full model R^2^ was 0.715, the change in R^2^ between full and reduced models was 0.077, F_df=3_ = 130.651. For DV, full model R^2^ was 0.715, the change in R^2^ between full and reduced models was 0.194, F_df=3_ = 340.634. For AP, full model R^2^ was 0.610, the change in R^2^ between full and reduced models was 0.470, F_df=3_ = 967.683.

**Fig. S3.**
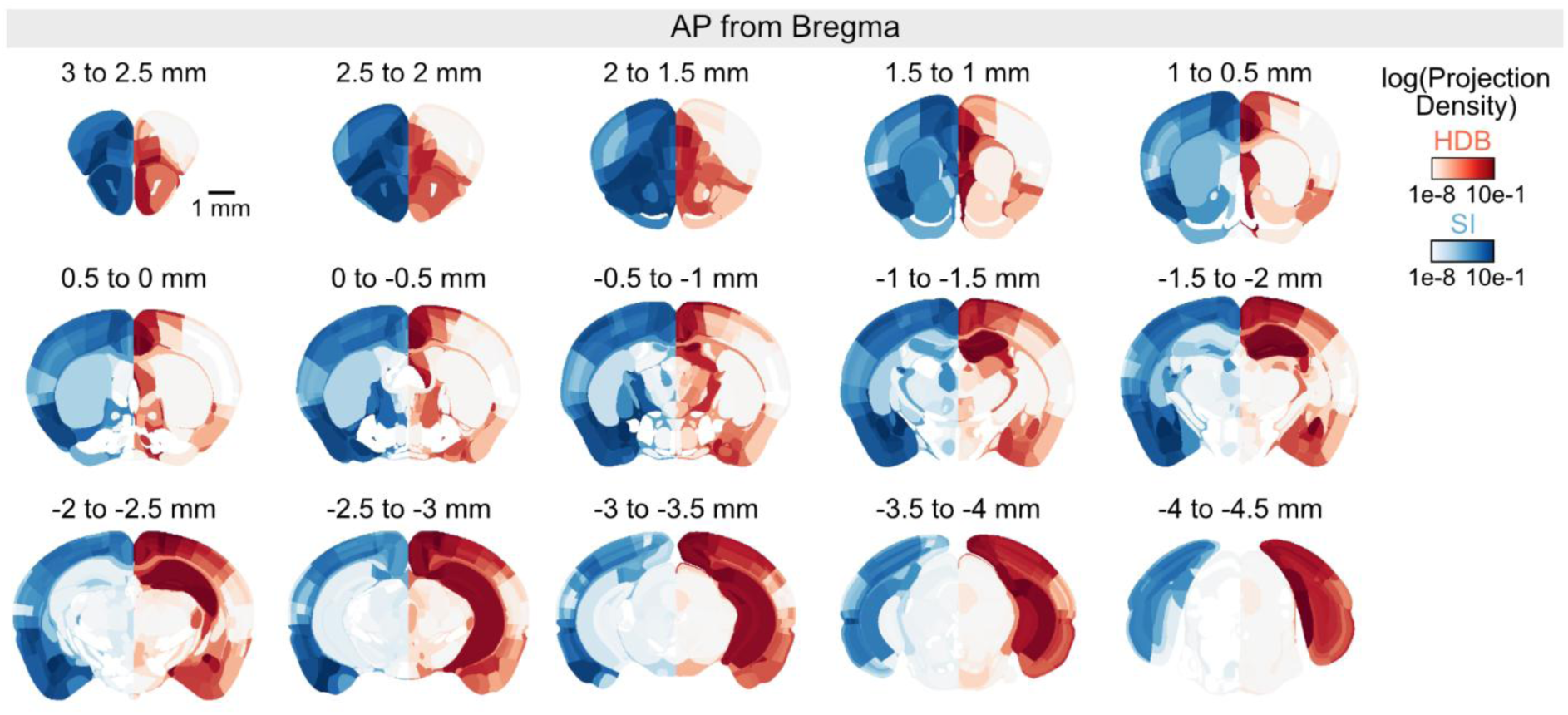
Additional quantification of projection density. Average cholinergic-projection density per cerebrum region from SI (left hemisphere, blue, n = 4) and HDB (right hemisphere, red, n = 3), for different 0.5-mm coronal volumes.

**Fig. S4.**
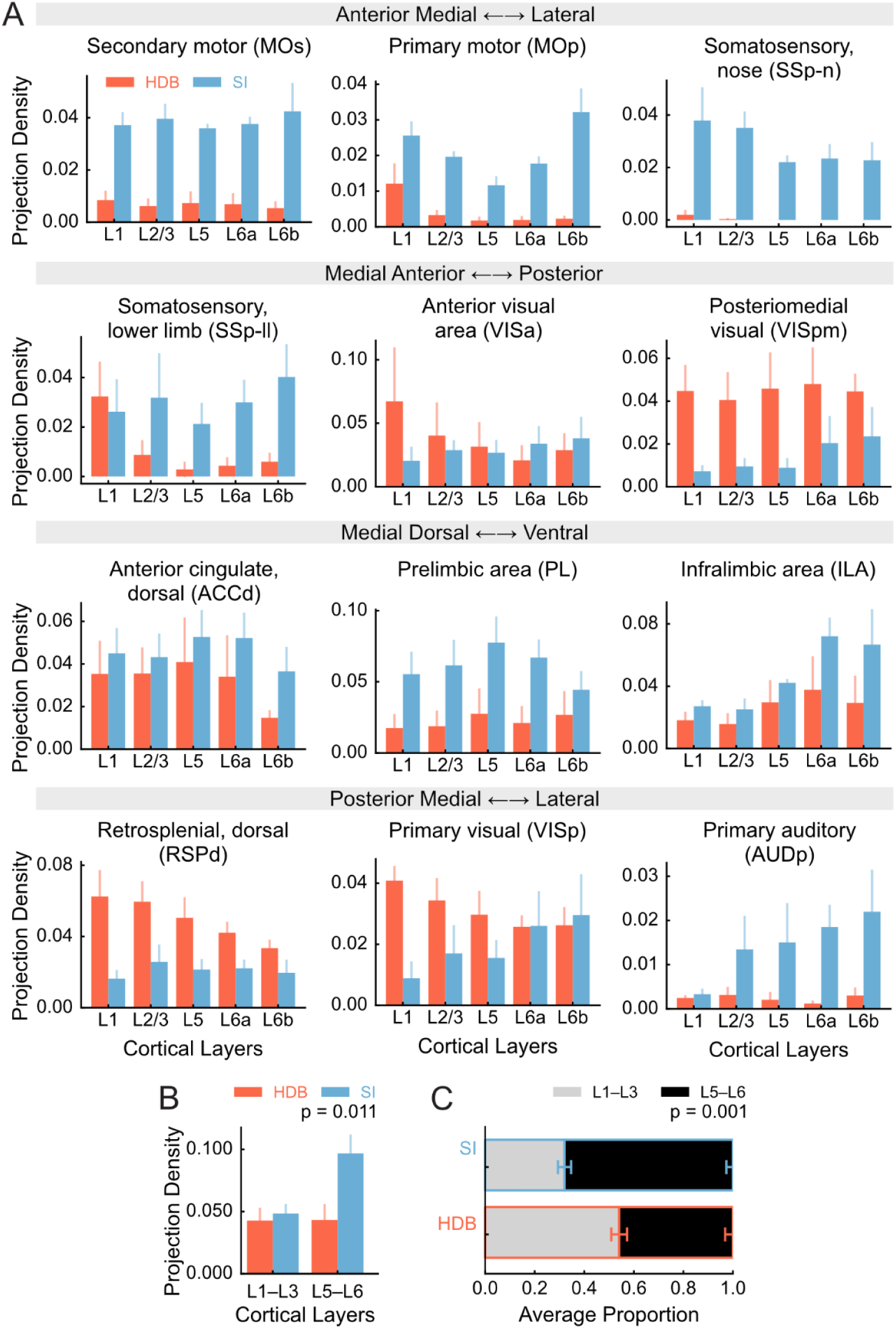
BF subnuclei project to distinct cortical layers in an area-dependent manner. (A) Average projection density from HDB (n = 3) and SI (n = 4) by target cortical layer, for example regions. Error bars: ± S.E.M. across mice. (B) Average projection density to superficial and deep layers combined across all cortical regions. Error bars: ± S.E.M. across mice. The printed p-value is for the interaction term of a mixed two-way ANOVA with factors injection location and cortical layer. (C) Proportion of projection density to superficial and deep layers, relative to the total input to each cortical region, averaged over mice and regions. Error bars: ± S.E.M. across mice. The printed p-value is for the interaction term of a mixed two-way ANOVA with factors injection location and cortical layer.

**Fig. S5.**
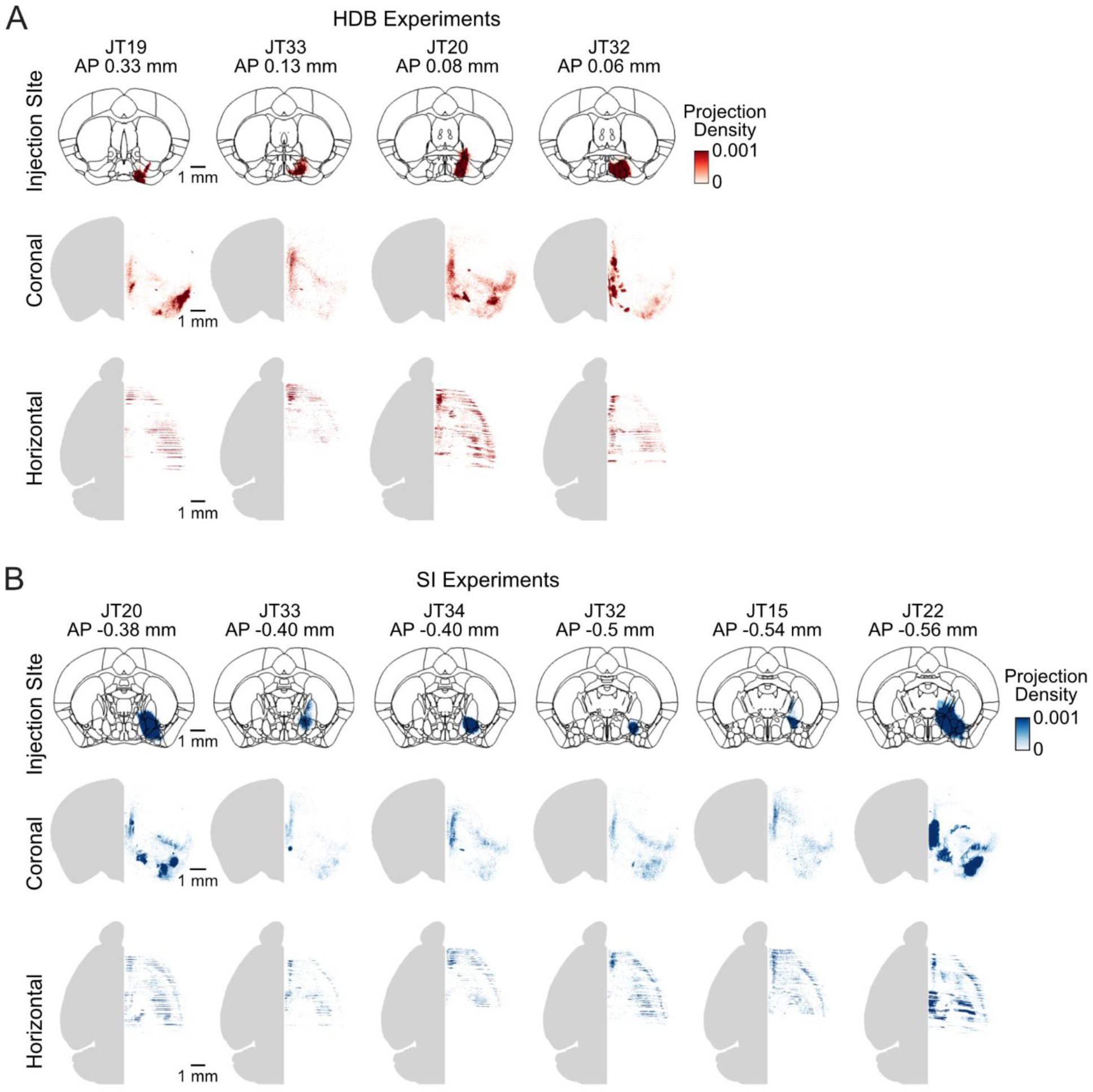
Injection site and input puncta from individual retrograde tracing experiments. (A) Top, injection locations for individual mice in the HDB dataset by animal ID. Bottom, average HDB-input puncta density per voxel, projected onto the coronal and horizontal planes for each brain. (B) Same as A, for the SI dataset.

**Fig. S6.**
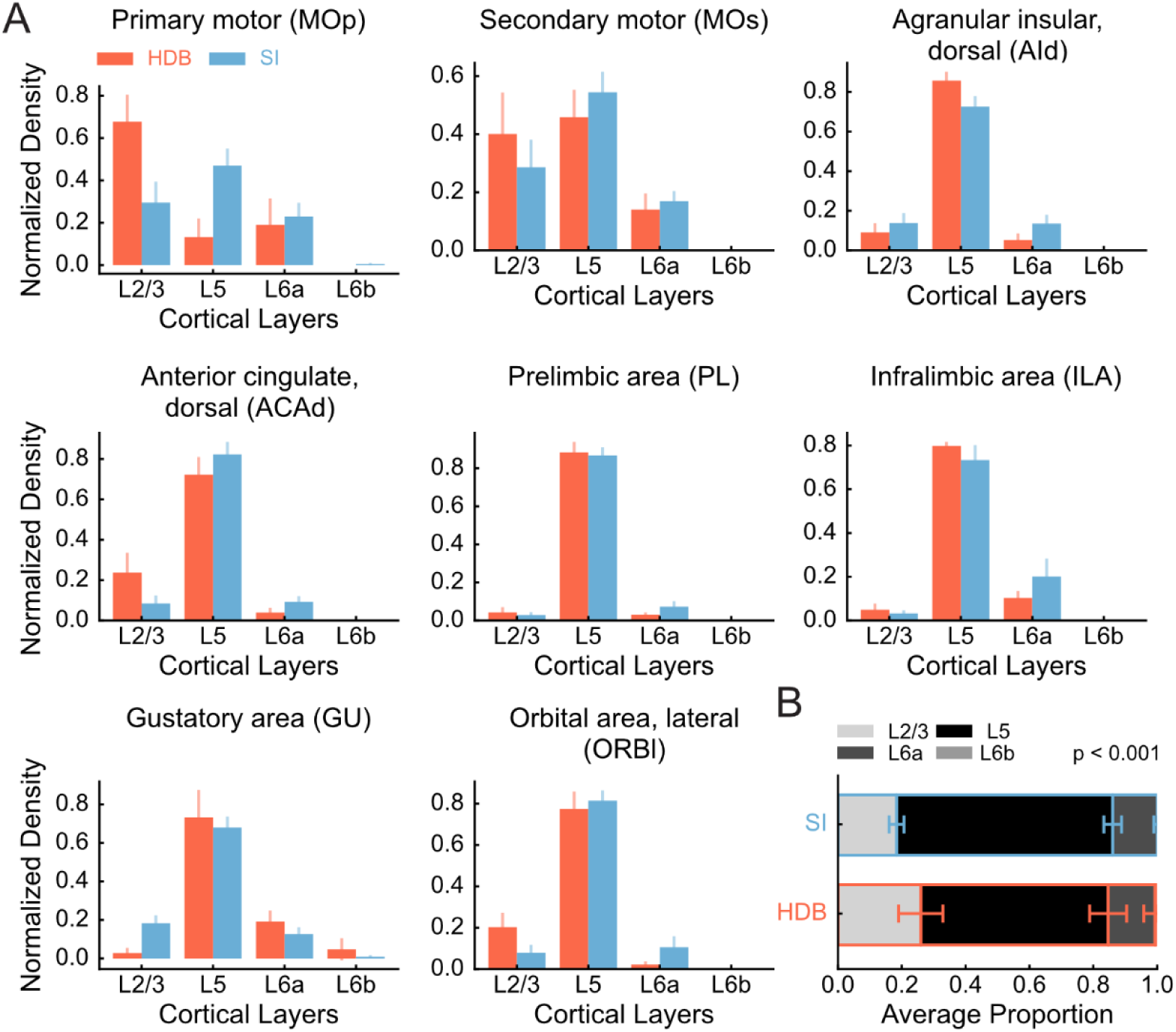
Both BF subnuclei primarily receive cortical input from layer 5. (A) Average density of puncta projecting to each subnucleus from different cortical layers, normalized to the total puncta count for each region, for example regions. Error bars: ± S.E.M. across mice (HDB: n = 4; SI: n = 6). (B) Average density of puncta projecting to each subnucleus from each layer, averaged across all cortical regions. Error bars: ± S.E.M. across mice. The printed p-value is for the interaction term of a two-way ANOVA with factors injection location and cortical layer.

**Fig. S7.**
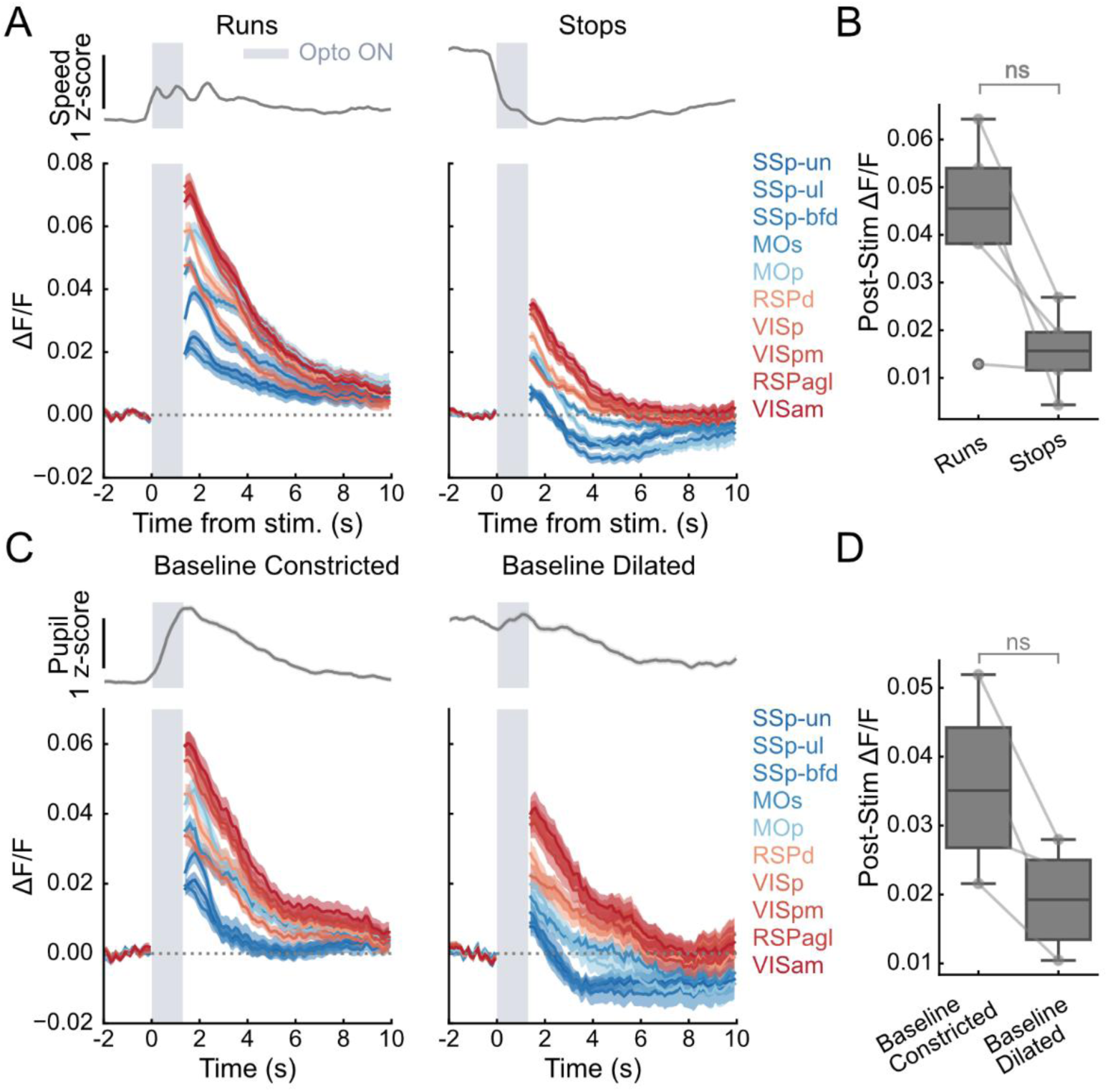
Behavioral state slightly modulates the effect of mPFC→HDB but does not affect our main conclusions. (A) We divided trials based on whether mice went from stationary to moving with stimulation (‘Runs’, n = 160/365 trials) or from running to stationary (‘Stops’, n = 205/365 trials) (see STAR methods). Top, average z-scored speed across the trial groups. Bottom, stimulation-aligned average traces for dorsal cortical regions, for each running condition. Error shades: ± S.E.M. across trials. (B) Box plots showing the distribution of responses per mouse (n = 5, average for 1 s following stimulation). p = 0.06 from a Wilcoxon signed-rank test. Lines: individual mice. Note that, while the magnitude of response decreases with ‘Stops’, it follows the same overall pattern. (C) Same as A, for pupil diameter. In this case, we divided trials in which pupils were constricted during the pre-stimulation baseline (‘Baseline Constricted’, n = 114/165trials), or dilated (‘Baseline Dilated’, n = 51/165trials). (D) Same as B, for pupil diameter. p = 0.13 from a Wilcoxon signed-rank test. Note that n = 4 for this analysis, since one mouse did not have eye tracking data.

**Table S1.** Brain region acronym definitions, average HDB and SI projection density, and average input density to HDB and SI.

| Acronym | Full name | Avg. HDB projection density | S.E.M. HDB projection density | Avg. SI proj. density | S.E.M. SI proj. density | Avg. input density to HDB | S.E.M. input density to HDB | Avg. input density to SI | S.E.M. input density to SI |
|---|---|---|---|---|---|---|---|---|---|
| FRP | Frontal pole, cerebral cortex | 9.42E-04 | 8.15E-04 | 5.07E-02 | 1.35E-02 | 4.73E-03 | 5.38E-03 | 5.14E-03 | 4.48E-03 |
| MOp | Primary motor area | 3.51E-03 | 1.54E-03 | 1.77E-02 | 1.68E-03 | 2.77E-03 | 1.64E-03 | 5.52E-03 | 2.38E-03 |
| MOs | Secondary motor area | 7.11E-03 | 3.74E-03 | 3.76E-02 | 2.57E-03 | 9.12E-03 | 7.43E-03 | 2.12E-02 | 8.05E-03 |
| SSp-n | Primary somatosensory area, nose | 3.46E-04 | 3.61E-04 | 2.77E-02 | 2.96E-03 | 1.44E-04 | 8.80E-05 | 1.30E-04 | 5.61E-05 |
| SSp-bfd | Primary somatosensory area, barrel field | 5.67E-03 | 3.72E-03 | 3.46E-02 | 9.62E-03 | 2.88E-04 | 1.72E-04 | 8.72E-05 | 5.57E-05 |
| SSp-ll | Primary somatosensory area, lower limb | 8.65E-03 | 3.94E-03 | 2.76E-02 | 1.16E-02 | 1.96E-04 | 1.98E-04 | 6.30E-04 | 3.85E-04 |
| SSp-m | Primary somatosensory area, mouth | 3.22E-05 | 2.27E-05 | 1.17E-02 | 6.02E-03 | 6.71E-04 | 6.95E-04 | 3.39E-04 | 1.82E-04 |
| SSp-ul | Primary somatosensory area, upper limb | 1.82E-03 | 1.05E-03 | 2.31E-02 | 4.03E-03 | 1.17E-03 | 9.36E-04 | 1.90E-03 | 1.27E-03 |
| SSp-tr | Primary somatosensory area, trunk | 2.28E-02 | 1.48E-02 | 2.57E-02 | 6.33E-03 | 3.18E-04 | 2.91E-04 | 2.57E-04 | 1.51E-04 |
| SSp-un | Primary somatosensory area, unassigned | 2.98E-03 | 1.84E-03 | 2.80E-02 | 2.66E-03 | 1.34E-04 | 8.83E-05 | 1.07E-04 | 7.76E-05 |
| SSs | Supplemental somatosensory area | 6.34E-04 | 3.61E-04 | 7.94E-03 | 1.83E-03 | 1.26E-03 | 6.61E-04 | 1.29E-03 | 4.45E-04 |
| GU | Gustatory areas | 7.26E-05 | 5.95E-05 | 6.41E-03 | 3.04E-03 | 3.20E-03 | 1.48E-03 | 9.01E-03 | 3.30E-03 |
| VISC | Visceral area | 1.82E-05 | 8.56E-06 | 1.21E-02 | 5.97E-03 | 4.59E-03 | 3.74E-03 | 7.21E-03 | 2.28E-03 |
| AUDd | Dorsal auditory area | 2.37E-03 | 1.36E-03 | 1.35E-02 | 6.64E-03 | 0.00E+00 | 0.00E+00 | 7.73E-05 | 8.47E-05 |
| AUDp | Primary auditory area | 1.19E-03 | 6.20E-04 | 7.31E-03 | 5.60E-03 | 3.58E-05 | 4.13E-05 | 2.67E-05 | 2.58E-05 |
| AUDpo | Posterior auditory area | 5.66E-03 | 2.43E-03 | 1.22E-02 | 7.48E-03 | 0.00E+00 | 0.00E+00 | 0.00E+00 | 0.00E+00 |
| AUDv | Ventral auditory area | 3.28E-04 | 2.50E-04 | 3.55E-03 | 1.66E-03 | 3.95E-04 | 2.63E-04 | 4.10E-04 | 3.56E-04 |
| VISal | Anterolateral visual area | 1.89E-02 | 1.76E-03 | 2.81E-02 | 1.09E-02 | 2.83E-05 | 3.26E-05 | 1.19E-05 | 8.31E-06 |
| VISam | Anteromedial visual area | 6.69E-02 | 3.31E-02 | 2.31E-02 | 1.35E-02 | 3.70E-05 | 2.47E-05 | 2.79E-04 | 2.20E-04 |
| VISl | Lateral visual area | 2.92E-02 | 4.51E-03 | 2.33E-02 | 1.17E-02 | 0.00E+00 | 0.00E+00 | 0.00E+00 | 0.00E+00 |
| VISp | Primary visual area | 3.40E-02 | 4.77E-03 | 1.79E-02 | 7.85E-03 | 3.58E-05 | 4.13E-05 | 5.22E-05 | 5.72E-05 |
| VISpl | Posterolateral visual area | 3.51E-02 | 9.52E-03 | 1.15E-02 | 5.98E-03 | 0.00E+00 | 0.00E+00 | 0.00E+00 | 0.00E+00 |
| VISpm | posteromedial visual area | 4.44E-02 | 1.33E-02 | 1.12E-02 | 4.11E-03 | 0.00E+00 | 0.00E+00 | 1.84E-05 | 1.42E-05 |
| VISli | Laterointermediate area | 2.00E-02 | 3.89E-03 | 1.90E-02 | 1.03E-02 | 0.00E+00 | 0.00E+00 | 0.00E+00 | 0.00E+00 |
| VISpor | Postrhinal area | 1.11E-02 | 2.57E-03 | 1.47E-02 | 5.55E-03 | 0.00E+00 | 0.00E+00 | 0.00E+00 | 0.00E+00 |
| ACAd | Anterior cingulate area, dorsal part | 3.70E-02 | 1.77E-02 | 4.91E-02 | 1.12E-02 | 6.99E-03 | 3.44E-03 | 1.09E-02 | 7.05E-03 |
| ACAv | Anterior cingulate area, ventral part | 8.30E-02 | 2.11E-02 | 2.17E-02 | 6.83E-03 | 1.04E-02 | 6.78E-03 | 1.01E-02 | 8.26E-03 |
| PL | Prelimbic area | 2.25E-02 | 1.37E-02 | 6.76E-02 | 1.53E-02 | 4.40E-02 | 3.19E-02 | 2.66E-02 | 1.10E-02 |
| ILA | Infralimbic area | 2.71E-02 | 1.28E-02 | 4.41E-02 | 2.75E-03 | 2.67E-02 | 4.34E-03 | 3.00E-02 | 1.14E-02 |
| ORBl | Orbital area, lateral part | 6.81E-03 | 4.08E-03 | 8.26E-02 | 1.17E-02 | 1.31E-02 | 9.39E-03 | 9.36E-03 | 4.42E-03 |
| ORBm | Orbital area, medial part | 1.93E-02 | 1.09E-02 | 3.58E-02 | 8.39E-03 | 4.26E-02 | 3.23E-02 | 2.55E-02 | 1.68E-02 |
| ORBvl | Orbital area, ventrolateral part | 2.52E-02 | 1.46E-02 | 4.53E-02 | 1.26E-02 | 2.54E-02 | 2.19E-02 | 7.10E-03 | 4.33E-03 |
| AId | Agranular insular area, dorsal part | 7.32E-04 | 4.49E-04 | 3.78E-02 | 1.32E-02 | 2.26E-02 | 1.24E-02 | 5.87E-02 | 1.27E-02 |
| AIp | Agranular insular area, posterior part | 9.20E-04 | 6.45E-04 | 3.80E-02 | 1.33E-02 | 1.71E-02 | 8.82E-03 | 2.10E-02 | 3.44E-03 |
| AIv | Agranular insular area, ventral part | 9.88E-03 | 7.66E-03 | 7.39E-02 | 2.27E-02 | 2.00E-02 | 9.26E-03 | 1.65E-02 | 5.42E-03 |
| RSPagl | Retrosplenial area, lateral agranular part | 6.37E-02 | 1.55E-02 | 1.86E-02 | 4.74E-03 | 3.60E-04 | 1.57E-04 | 4.59E-04 | 1.96E-04 |
| RSPd | Retrosplenial area, dorsal part | 5.36E-02 | 8.47E-03 | 2.16E-02 | 6.14E-03 | 7.05E-04 | 2.57E-04 | 1.84E-04 | 7.29E-05 |
| RSPv | Retrosplenial area, ventral part | 5.56E-02 | 4.99E-03 | 1.54E-02 | 3.96E-03 | 1.86E-03 | 1.63E-03 | 5.43E-04 | 4.37E-04 |
| VISa | Anterior area | 3.86E-02 | 2.44E-02 | 2.79E-02 | 9.11E-03 | 7.53E-05 | 6.25E-05 | 2.28E-05 | 1.46E-05 |
| VISrl | Rostrolateral visual area | 2.03E-02 | 1.26E-02 | 2.66E-02 | 6.91E-03 | 0.00E+00 | 0.00E+00 | 5.49E-06 | 6.02E-06 |
| TEa | Temporal association areas | 2.14E-03 | 6.11E-04 | 8.93E-03 | 3.93E-03 | 3.55E-03 | 2.79E-03 | 1.09E-03 | 4.63E-04 |
| PERI | Perirhinal area | 8.59E-03 | 3.58E-03 | 4.22E-02 | 3.57E-03 | 1.63E-03 | 1.17E-03 | 6.30E-04 | 4.25E-04 |
| ECT | Ectorhinal area | 2.85E-03 | 4.86E-04 | 3.71E-02 | 1.17E-02 | 5.04E-03 | 5.00E-03 | 2.60E-03 | 1.79E-03 |
| MOB | Main olfactory bulb | 1.73E-02 | 7.94E-04 | 4.39E-02 | 1.85E-02 | 6.81E-04 | 4.37E-04 | 9.29E-05 | 9.82E-05 |
| AOB | Accessory olfactory bulb | 2.20E-02 | 8.75E-03 | 9.37E-03 | 5.93E-03 | 0.00E+00 | 0.00E+00 | 0.00E+00 | 0.00E+00 |
| AON | Anterior olfactory nucleus | 1.10E-02 | 3.69E-03 | 7.22E-02 | 2.22E-02 | 4.28E-02 | 1.66E-02 | 6.92E-03 | 3.89E-03 |
| TT | Taenia tecta | 2.05E-02 | 7.42E-03 | 4.85E-02 | 1.71E-02 | 1.69E-02 | 5.58E-03 | 5.17E-03 | 3.29E-03 |
| DP | Dorsal peduncular area | 3.98E-02 | 1.82E-02 | 5.08E-02 | 1.36E-02 | 1.30E-02 | 4.45E-03 | 1.18E-02 | 9.14E-03 |
| PIR | Piriform area | 3.67E-03 | 1.91E-03 | 5.40E-02 | 2.40E-02 | 9.95E-02 | 7.80E-02 | 1.76E-02 | 7.74E-03 |
| NLOT | Nucleus of the lateral olfactory tract | 1.69E-02 | 9.84E-03 | 4.86E-02 | 2.90E-02 | 1.11E-02 | 5.08E-03 | 7.05E-03 | 3.84E-03 |
| COAa | Cortical amygdalar area, anterior part | 1.86E-02 | 1.10E-02 | 4.90E-02 | 2.71E-02 | 1.93E-02 | 7.78E-03 | 1.81E-02 | 5.92E-03 |
| COAp | Cortical amygdalar area, posterior part | 1.27E-02 | 8.45E-03 | 2.68E-02 | 1.10E-02 | 5.44E-03 | 3.54E-03 | 1.28E-02 | 5.79E-03 |
| PAA | Piriform-amygdalar area | 8.92E-03 | 5.37E-03 | 5.07E-02 | 2.69E-02 | 9.21E-03 | 1.87E-03 | 5.08E-03 | 1.48E-03 |
| TR | Postpiriform transition area | 1.15E-02 | 8.20E-03 | 7.61E-02 | 2.41E-02 | 2.39E-02 | 2.25E-02 | 1.84E-03 | 9.47E-04 |
| CA1 | Field CA1 | 6.09E-02 | 3.50E-02 | 2.08E-03 | 8.60E-04 | 2.11E-03 | 1.15E-03 | 7.60E-04 | 4.39E-04 |
| CA2 | Field CA2 | 6.13E-02 | 3.58E-02 | 2.11E-03 | 1.83E-03 | 1.91E-05 | 2.21E-05 | 0.00E+00 | 0.00E+00 |
| CA3 | Field CA3 | 6.69E-02 | 3.53E-02 | 2.31E-03 | 1.88E-03 | 3.84E-04 | 3.35E-04 | 5.07E-05 | 5.20E-05 |
| ENTl | Entorhinal area, lateral part | 1.38E-02 | 8.45E-03 | 2.41E-02 | 9.86E-03 | 3.67E-02 | 3.80E-02 | 1.62E-03 | 8.10E-04 |
| ENTm | Entorhinal area, medial part, dorsal zone | 1.80E-02 | 1.50E-02 | 2.21E-03 | 8.84E-04 | 6.69E-04 | 7.73E-04 | 0.00E+00 | 0.00E+00 |
| PAR | Parasubiculum | 7.96E-02 | 3.45E-02 | 2.07E-03 | 1.14E-03 | 0.00E+00 | 0.00E+00 | 0.00E+00 | 0.00E+00 |
| POST | Postsubiculum | 6.88E-02 | 2.14E-02 | 1.16E-02 | 2.48E-03 | 0.00E+00 | 0.00E+00 | 0.00E+00 | 0.00E+00 |
| PRE | Presubiculum | 1.16E-01 | 3.30E-02 | 5.26E-03 | 2.21E-03 | 0.00E+00 | 0.00E+00 | 0.00E+00 | 0.00E+00 |
| SUB | Subiculum | 7.92E-02 | 2.97E-02 | 1.31E-02 | 4.71E-03 | 2.72E-04 | 2.88E-04 | 4.41E-04 | 3.16E-04 |
| ProS | Prosubiculum | 6.62E-02 | 3.28E-02 | 1.73E-02 | 6.36E-03 | 2.97E-04 | 1.20E-04 | 8.44E-05 | 6.09E-05 |
| HATA | Hippocampo-amygdalar transition area | 8.26E-03 | 8.12E-03 | 5.78E-04 | 2.36E-04 | 3.35E-04 | 1.29E-04 | 1.42E-04 | 7.32E-05 |
| APr | Area prostriata | 7.26E-02 | 1.33E-02 | 1.46E-03 | 7.89E-04 | 3.35E-04 | 1.29E-04 | 1.42E-04 | 7.32E-05 |
| CLA | Claustrum | 1.40E-02 | 7.95E-03 | 6.75E-02 | 9.95E-03 | 1.03E-03 | 7.06E-04 | 2.67E-04 | 1.03E-04 |
| LA | Lateral amygdalar nucleus | 4.27E-03 | 3.33E-03 | 3.31E-02 | 9.37E-03 | 2.69E-03 | 1.35E-03 | 4.87E-03 | 2.48E-03 |
| BLA | Basolateral amygdalar nucleus | 3.62E-02 | 1.72E-02 | 1.48E-01 | 2.36E-02 | 5.24E-02 | 2.71E-02 | 1.00E-01 | 3.63E-02 |
| BMA | Basomedial amygdalar nucleus | 1.86E-02 | 8.92E-03 | 7.83E-02 | 2.54E-02 | 1.92E-02 | 5.16E-03 | 4.71E-02 | 1.62E-02 |
| PA | Posterior amygdalar nucleus | 2.12E-02 | 1.73E-02 | 4.05E-02 | 1.47E-02 | 2.23E-02 | 6.40E-03 | 5.73E-02 | 1.92E-02 |
| CP | Caudoputamen | 2.00E-04 | 7.65E-05 | 4.96E-03 | 7.25E-04 | 3.15E-02 | 6.55E-03 | 6.96E-02 | 1.87E-02 |
| ACB | Nucleus accumbens | 2.44E-03 | 1.64E-03 | 1.32E-02 | 4.24E-03 | 7.36E-02 | 3.64E-02 | 8.39E-02 | 1.43E-02 |
| FS | Fundus of striatum | 5.71E-04 | 4.69E-04 | 5.30E-02 | 1.07E-02 | 7.39E-02 | 3.64E-02 | 8.56E-02 | 1.41E-02 |
| OT | Olfactory tubercle | 1.66E-03 | 6.18E-04 | 7.81E-03 | 3.03E-03 | 2.14E-03 | 8.76E-04 | 2.07E-03 | 5.58E-04 |
| LSc | Lateral septal nucleus, caudal (caudodorsal) part | 6.31E-03 | 3.55E-03 | 1.94E-04 | 1.93E-04 | 4.65E-03 | 2.19E-03 | 2.13E-03 | 5.79E-04 |
| LSr | Lateral septal nucleus, rostral (rostroventral) part | 2.44E-02 | 1.20E-02 | 8.43E-04 | 4.77E-04 | 3.01E-02 | 2.90E-02 | 2.22E-03 | 5.88E-04 |
| LSv | Lateral septal nucleus, ventral part | 2.98E-03 | 2.83E-03 | 9.86E-05 | 7.76E-05 | 3.07E-02 | 2.95E-02 | 2.22E-03 | 5.85E-04 |
| SF | Septofimbrial nucleus | 5.08E-02 | 3.50E-02 | 1.28E-03 | 1.36E-03 | 4.00E-03 | 1.79E-03 | 2.08E-03 | 5.54E-04 |
| SH | Septohippocampal nucleus | 2.52E-02 | 1.27E-02 | 4.08E-03 | 1.67E-03 | 4.22E-03 | 1.91E-03 | 2.10E-03 | 5.63E-04 |
| AAA | Anterior amygdalar area | 1.17E-02 | 4.81E-03 | 8.62E-02 | 3.95E-02 | 3.28E-02 | 2.92E-02 | 8.14E-03 | 4.25E-03 |
| BA | Bed nucleus of the accessory olfactory tract | 3.18E-02 | 2.18E-02 | 1.61E-02 | 1.28E-02 | 3.28E-02 | 2.92E-02 | 8.15E-03 | 4.24E-03 |
| CEA | Central amygdalar nucleus | 6.39E-03 | 3.90E-03 | 7.81E-02 | 1.80E-02 | 1.68E-03 | 1.38E-03 | 4.23E-02 | 3.59E-02 |
| IA | Intercalated amygdalar nucleus | 1.21E-02 | 5.46E-03 | 7.27E-02 | 2.29E-02 | 2.20E-03 | 1.64E-03 | 4.40E-02 | 3.59E-02 |
| MEA | Medial amygdalar nucleus | 1.83E-02 | 1.14E-02 | 2.55E-02 | 1.03E-02 | 4.62E-03 | 2.36E-03 | 1.33E-02 | 3.37E-03 |
| GPe | Globus pallidus, external segment | 2.57E-05 | 1.69E-05 | 1.20E-02 | 6.69E-03 | 4.82E-03 | 2.41E-03 | 1.46E-02 | 3.74E-03 |
| GPi | Globus pallidus, internal segment | 2.72E-05 | 1.68E-05 | 5.21E-03 | 2.60E-03 | 4.85E-03 | 2.44E-03 | 1.74E-02 | 4.88E-03 |
| TRS | Triangular nucleus of septum | 0.00E+00 | 0.00E+00 | 0.00E+00 | 0.00E+00 | 5.40E-03 | 2.97E-03 | 1.74E-02 | 4.88E-03 |
| BST | Bed nuclei of the stria terminalis | 8.91E-03 | 3.48E-03 | 2.38E-02 | 6.70E-03 | 2.27E-03 | 1.14E-03 | 4.14E-03 | 3.03E-03 |
| BAC | Bed nucleus of the anterior commissure | 9.67E-03 | 4.98E-03 | 1.07E-02 | 7.07E-03 | 2.27E-03 | 1.14E-03 | 4.14E-03 | 3.03E-03 |
| VAL | Ventral anterior-lateral complex of the thalamus | 3.10E-04 | 8.03E-05 | 2.13E-03 | 1.77E-03 | N/A | N/A | N/A | N/A |
| VM | Ventral medial nucleus of the thalamus | 2.79E-04 | 1.45E-04 | 1.58E-03 | 1.07E-03 | N/A | N/A | N/A | N/A |
| VPL | Ventral posterolateral nucleus of the thalamus | 3.17E-04 | 9.89E-05 | 9.05E-04 | 6.38E-04 | N/A | N/A | N/A | N/A |
| VPLpc | Ventral posterolateral nucleus of the thalamus, parvicellular part | 1.54E-05 | 1.41E-05 | 3.92E-04 | 1.31E-04 | N/A | N/A | N/A | N/A |
| VPM | Ventral posteromedial nucleus of the thalamus | 2.14E-05 | 1.64E-05 | 5.04E-04 | 3.87E-04 | N/A | N/A | N/A | N/A |
| VPMpc | Ventral posteromedial nucleus of the thalamus, parvicellular part | 6.51E-05 | 5.02E-05 | 1.32E-03 | 6.45E-04 | N/A | N/A | N/A | N/A |
| PoT | Posterior triangular thalamic nucleus | 6.18E-06 | 4.31E-06 | 9.99E-05 | 6.88E-05 | N/A | N/A | N/A | N/A |
| SPA | Subparafascicular area | 1.35E-04 | 7.58E-05 | 2.38E-03 | 1.73E-03 | N/A | N/A | N/A | N/A |
| PP | Peripeduncular nucleus | 5.10E-04 | 6.04E-04 | 3.00E-05 | 3.47E-05 | N/A | N/A | N/A | N/A |
| MG | Medial geniculate complex | 2.17E-04 | 1.24E-04 | 1.56E-04 | 8.14E-05 | N/A | N/A | N/A | N/A |
| LGd | Dorsal part of the lateral geniculate complex | 7.37E-04 | 6.08E-04 | 9.27E-05 | 5.59E-05 | N/A | N/A | N/A | N/A |
| LP | Lateral posterior nucleus of the thalamus | 1.69E-03 | 7.07E-04 | 1.19E-04 | 1.10E-04 | N/A | N/A | N/A | N/A |
| PO | Posterior complex of the thalamus | 1.13E-04 | 5.19E-05 | 4.37E-04 | 2.59E-04 | N/A | N/A | N/A | N/A |
| POL | Posterior limiting nucleus of the thalamus | 2.55E-04 | 1.44E-04 | 1.20E-04 | 8.24E-05 | N/A | N/A | N/A | N/A |
| SGN | Suprageniculate nucleus | 3.46E-04 | 2.70E-04 | 2.03E-05 | 1.40E-05 | N/A | N/A | N/A | N/A |
| AV | Anteroventral nucleus of thalamus | 4.51E-03 | 6.72E-04 | 7.53E-04 | 4.87E-04 | N/A | N/A | N/A | N/A |
| AM | Anteromedial nucleus | 5.62E-03 | 8.01E-04 | 1.36E-03 | 7.42E-04 | N/A | N/A | N/A | N/A |
| AD | Anterodorsal nucleus | 4.15E-03 | 2.29E-03 | 1.57E-04 | 1.45E-04 | N/A | N/A | N/A | N/A |
| IAM | Interanteromedial nucleus of the thalamus | 2.34E-03 | 8.60E-04 | 4.28E-03 | 3.09E-03 | N/A | N/A | N/A | N/A |
| IAD | Interanterodorsal nucleus of the thalamus | 1.58E-03 | 6.46E-04 | 6.79E-04 | 4.17E-04 | N/A | N/A | N/A | N/A |
| LD | Lateral dorsal nucleus of thalamus | 5.61E-03 | 1.15E-03 | 9.56E-05 | 7.89E-05 | N/A | N/A | N/A | N/A |
| IMD | Intermediodorsal nucleus of the thalamus | 5.77E-05 | 4.10E-05 | 3.57E-03 | 2.07E-03 | N/A | N/A | N/A | N/A |
| MD | Mediodorsal nucleus of thalamus | 1.03E-03 | 4.41E-04 | 7.21E-03 | 5.47E-03 | N/A | N/A | N/A | N/A |
| SMT | Submedial nucleus of the thalamus | 2.05E-03 | 1.57E-03 | 3.39E-03 | 3.19E-04 | N/A | N/A | N/A | N/A |
| PR | Perireunensis nucleus | 3.82E-03 | 2.24E-03 | 8.90E-04 | 5.18E-04 | N/A | N/A | N/A | N/A |
| PVT | Paraventricular nucleus of the thalamus | 4.16E-04 | 4.21E-04 | 8.73E-04 | 3.29E-04 | N/A | N/A | N/A | N/A |
| PT | Parataenial nucleus | 2.71E-03 | 2.68E-03 | 1.11E-03 | 5.65E-04 | N/A | N/A | N/A | N/A |
| RE | Nucleus of reuniens | 3.02E-03 | 1.57E-03 | 9.39E-04 | 3.56E-04 | N/A | N/A | N/A | N/A |
| Xi | Xiphoid thalamic nucleus | 2.67E-03 | 2.33E-03 | 1.37E-03 | 8.11E-04 | N/A | N/A | N/A | N/A |
| RH | Rhomboid nucleus | 3.72E-03 | 2.06E-03 | 7.24E-04 | 1.84E-04 | N/A | N/A | N/A | N/A |
| CM | Central medial nucleus of the thalamus | 1.69E-04 | 1.20E-04 | 2.03E-03 | 7.39E-04 | N/A | N/A | N/A | N/A |
| PCN | Paracentral nucleus | 3.79E-05 | 2.89E-06 | 1.15E-03 | 7.58E-04 | N/A | N/A | N/A | N/A |
| CL | Central lateral nucleus of the thalamus | 1.04E-03 | 2.69E-04 | 6.78E-04 | 2.70E-04 | N/A | N/A | N/A | N/A |
| PF | Parafascicular nucleus | 1.27E-04 | 7.30E-05 | 5.13E-04 | 3.71E-04 | N/A | N/A | N/A | N/A |
| PIL | Posterior intralaminar thalamic nucleus | 2.77E-04 | 1.69E-04 | 1.61E-04 | 1.85E-04 | N/A | N/A | N/A | N/A |
| RT | Reticular nucleus of the thalamus | 1.68E-02 | 4.14E-03 | 2.19E-02 | 1.10E-02 | N/A | N/A | N/A | N/A |
| IGL | Intergeniculate leaflet of the lateral geniculate complex | 3.43E-03 | 2.88E-03 | 1.67E-04 | 9.04E-05 | N/A | N/A | N/A | N/A |
| IntG | Intermediate geniculate nucleus | 1.06E-03 | 8.69E-04 | 3.00E-05 | 3.46E-05 | N/A | N/A | N/A | N/A |
| LGv | Ventral part of the lateral geniculate complex | 1.06E-02 | 4.16E-03 | 3.60E-04 | 2.69E-04 | N/A | N/A | N/A | N/A |
| SubG | Subgeniculate nucleus | 3.85E-04 | 4.69E-04 | 3.86E-04 | 2.88E-04 | N/A | N/A | N/A | N/A |
| MH | Medial habenula | 3.34E-04 | 2.49E-04 | 5.95E-05 | 5.29E-05 | N/A | N/A | N/A | N/A |
| LH | Lateral habenula | 3.18E-03 | 2.64E-03 | 1.01E-02 | 8.42E-03 | N/A | N/A | N/A | N/A |
| SO | Supraoptic nucleus | 2.04E-02 | 8.06E-03 | 6.27E-03 | 5.03E-03 | N/A | N/A | N/A | N/A |
| ASO | Accessory supraoptic group | 2.08E-03 | 1.24E-03 | 7.97E-07 | 9.20E-07 | N/A | N/A | N/A | N/A |
| PVH | Paraventricular hypothalamic nucleus | 2.58E-05 | 2.78E-05 | 6.36E-04 | 4.07E-04 | N/A | N/A | N/A | N/A |
| PVa | Periventricular hypothalamic nucleus, anterior part | 0.00E+00 | 0.00E+00 | 3.69E-05 | 4.27E-05 | N/A | N/A | N/A | N/A |
| PVi | Periventricular hypothalamic nucleus, intermediate part | 1.26E-05 | 1.55E-05 | 8.35E-05 | 8.94E-05 | N/A | N/A | N/A | N/A |
| ARH | Arcuate hypothalamic nucleus | 4.53E-04 | 2.92E-04 | 4.18E-05 | 2.06E-05 | N/A | N/A | N/A | N/A |
| ADP | Anterodorsal preoptic nucleus | 3.93E-03 | 3.59E-03 | 5.74E-05 | 5.19E-05 | N/A | N/A | N/A | N/A |
| AVP | Anteroventral preoptic nucleus | 2.05E-02 | 5.24E-03 | 4.27E-04 | 4.25E-04 | N/A | N/A | N/A | N/A |
| AVPV | Anteroventral periventricular nucleus | 2.48E-02 | 2.16E-02 | 1.66E-05 | 1.01E-05 | N/A | N/A | N/A | N/A |
| DMH | Dorsomedial nucleus of the hypothalamus | 1.62E-04 | 1.62E-04 | 4.28E-05 | 4.94E-05 | N/A | N/A | N/A | N/A |
| OV | Vascular organ of the lamina terminalis | 1.26E-03 | 7.83E-04 | 6.37E-11 | 7.36E-11 | N/A | N/A | N/A | N/A |
| PD | Posterodorsal preoptic nucleus | 4.01E-05 | 2.67E-05 | 1.77E-03 | 1.88E-03 | N/A | N/A | N/A | N/A |
| PS | Parastrial nucleus | 1.81E-02 | 1.42E-02 | 5.33E-04 | 2.71E-04 | N/A | N/A | N/A | N/A |
| PVp | Periventricular hypothalamic nucleus, posterior part | 1.35E-04 | 7.09E-05 | 1.00E-04 | 1.10E-04 | N/A | N/A | N/A | N/A |
| PVpo | Periventricular hypothalamic nucleus, preoptic part | 4.69E-04 | 2.98E-04 | 3.25E-05 | 3.75E-05 | N/A | N/A | N/A | N/A |
| SBPV | Subparaventricular zone | 9.99E-07 | 1.22E-06 | 0.00E+00 | 0.00E+00 | N/A | N/A | N/A | N/A |
| SCH | Suprachiasmatic nucleus | 6.14E-04 | 7.52E-04 | 0.00E+00 | 0.00E+00 | N/A | N/A | N/A | N/A |
| SFO | Subfornical organ | 3.37E-03 | 3.45E-03 | 9.14E-06 | 6.42E-06 | N/A | N/A | N/A | N/A |
| VMPO | Ventromedial preoptic nucleus | 3.21E-04 | 1.19E-04 | 0.00E+00 | 0.00E+00 | N/A | N/A | N/A | N/A |
| VLPO | Ventrolateral preoptic nucleus | 2.74E-02 | 1.37E-02 | 6.13E-04 | 6.88E-04 | N/A | N/A | N/A | N/A |
| AHN | Anterior hypothalamic nucleus | 1.92E-03 | 1.02E-03 | 1.61E-04 | 6.37E-05 | N/A | N/A | N/A | N/A |
| MPN | Medial preoptic nucleus | 3.99E-04 | 1.28E-04 | 7.80E-05 | 5.23E-05 | N/A | N/A | N/A | N/A |
| PMd | Dorsal premammillary nucleus | 1.41E-03 | 8.16E-04 | 9.14E-07 | 1.06E-06 | N/A | N/A | N/A | N/A |
| PMv | Ventral premammillary nucleus | 4.97E-05 | 6.08E-05 | 1.12E-05 | 7.43E-06 | N/A | N/A | N/A | N/A |
| PVHd | Paraventricular hypothalamic nucleus, descending division | 1.71E-03 | 6.78E-04 | 1.62E-03 | 1.30E-03 | N/A | N/A | N/A | N/A |
| VMH | Ventromedial hypothalamic nucleus | 3.14E-04 | 1.29E-04 | 2.60E-03 | 2.21E-03 | N/A | N/A | N/A | N/A |
| PH | Posterior hypothalamic nucleus | 3.08E-03 | 1.94E-03 | 2.88E-04 | 2.81E-04 | N/A | N/A | N/A | N/A |
| LHA | Lateral hypothalamic area | 0.00E+00 | 0.00E+00 | 0.00E+00 | 0.00E+00 | N/A | N/A | N/A | N/A |
| LPO | Lateral preoptic area | 0.00E+00 | 0.00E+00 | 0.00E+00 | 0.00E+00 | N/A | N/A | N/A | N/A |
| PST | Preparasubthalamic nucleus | 0.00E+00 | 0.00E+00 | 0.00E+00 | 0.00E+00 | N/A | N/A | N/A | N/A |
| PSTN | Parasubthalamic nucleus | 0.00E+00 | 0.00E+00 | 0.00E+00 | 0.00E+00 | N/A | N/A | N/A | N/A |
| PeF | Perifornical nucleus | 0.00E+00 | 0.00E+00 | 0.00E+00 | 0.00E+00 | N/A | N/A | N/A | N/A |
| RCH | Retrochiasmatic area | 0.00E+00 | 0.00E+00 | 0.00E+00 | 0.00E+00 | N/A | N/A | N/A | N/A |
| STN | Subthalamic nucleus | 0.00E+00 | 0.00E+00 | 0.00E+00 | 0.00E+00 | N/A | N/A | N/A | N/A |
| TU | Tuberal nucleus | 0.00E+00 | 0.00E+00 | 0.00E+00 | 0.00E+00 | N/A | N/A | N/A | N/A |
| ZI | Zona incerta | 0.00E+00 | 0.00E+00 | 0.00E+00 | 0.00E+00 | N/A | N/A | N/A | N/A |
| ME | Median eminence | 7.05E-04 | 4.03E-04 | 2.80E-04 | 1.64E-04 | N/A | N/A | N/A | N/A |
| SCs | Superior colliculus, sensory related | 2.08E-03 | 1.30E-03 | 2.63E-04 | 1.16E-04 | N/A | N/A | N/A | N/A |
| IC | Inferior colliculus | 1.79E-07 | 9.62E-08 | 9.69E-06 | 6.18E-06 | N/A | N/A | N/A | N/A |
| NB | Nucleus of the brachium of the inferior colliculus | 6.72E-06 | 6.30E-06 | 1.22E-05 | 5.60E-06 | N/A | N/A | N/A | N/A |
| SAG | Nucleus sagulum | 0.00E+00 | 0.00E+00 | 7.67E-05 | 8.85E-05 | N/A | N/A | N/A | N/A |
| PBG | Parabigeminal nucleus | 2.74E-07 | 3.36E-07 | 1.83E-05 | 1.21E-05 | N/A | N/A | N/A | N/A |
| MEV | Midbrain trigeminal nucleus | 0.00E+00 | 0.00E+00 | 1.03E-04 | 1.19E-04 | N/A | N/A | N/A | N/A |
| SCO | Subcommissural organ | 3.47E-05 | 4.26E-05 | 0.00E+00 | 0.00E+00 | N/A | N/A | N/A | N/A |
| SNr | Substantia nigra, reticular part | 9.99E-06 | 3.38E-06 | 7.08E-04 | 4.55E-04 | N/A | N/A | N/A | N/A |
| VTA | Ventral tegmental area | 1.03E-03 | 1.04E-04 | 7.48E-04 | 3.65E-04 | N/A | N/A | N/A | N/A |
| PN | Paranigral nucleus | 3.28E-04 | 2.77E-04 | 7.71E-06 | 8.91E-06 | N/A | N/A | N/A | N/A |
| RR | Midbrain reticular nucleus, retrorubral area | 1.50E-04 | 1.80E-04 | 1.44E-04 | 1.53E-04 | N/A | N/A | N/A | N/A |
| MRN | Midbrain reticular nucleus | 7.01E-05 | 9.63E-06 | 3.84E-04 | 1.16E-04 | N/A | N/A | N/A | N/A |
| SCm | Superior colliculus, motor related | 6.87E-05 | 3.73E-05 | 3.99E-04 | 2.89E-04 | N/A | N/A | N/A | N/A |
| PAG | Periaqueductal gray | 8.77E-04 | 7.47E-04 | 1.70E-04 | 3.46E-05 | N/A | N/A | N/A | N/A |
| CUN | Cuneiform nucleus | 1.74E-05 | 1.04E-05 | 6.02E-05 | 3.18E-05 | N/A | N/A | N/A | N/A |
| RN | Red nucleus | 1.03E-04 | 7.36E-05 | 3.42E-04 | 1.57E-04 | N/A | N/A | N/A | N/A |
| III | Oculomotor nucleus | 6.02E-06 | 7.32E-06 | 1.58E-05 | 1.75E-05 | N/A | N/A | N/A | N/A |
| MA3 | Medial accesory oculomotor nucleus | 4.26E-05 | 5.22E-05 | 3.65E-04 | 4.14E-04 | N/A | N/A | N/A | N/A |
| EW | Edinger-Westphal nucleus | 1.86E-04 | 2.14E-04 | 3.44E-05 | 3.93E-05 | N/A | N/A | N/A | N/A |
| IV | Trochlear nucleus | 4.66E-04 | 5.71E-04 | 0.00E+00 | 0.00E+00 | N/A | N/A | N/A | N/A |
| Pa4 | Paratrochlear nucleus | 2.51E-05 | 3.07E-05 | 0.00E+00 | 0.00E+00 | N/A | N/A | N/A | N/A |
| VTN | Ventral tegmental nucleus | 3.06E-06 | 3.74E-06 | 0.00E+00 | 0.00E+00 | N/A | N/A | N/A | N/A |
| AT | Anterior tegmental nucleus | 0.00E+00 | 0.00E+00 | 0.00E+00 | 0.00E+00 | N/A | N/A | N/A | N/A |
| LT | Lateral terminal nucleus of the accessory optic tract | 2.70E-04 | 1.77E-04 | 8.20E-06 | 5.49E-06 | N/A | N/A | N/A | N/A |
| DT | Dorsal terminal nucleus of the accessory optic tract | 5.00E-04 | 6.07E-04 | 1.66E-03 | 1.05E-03 | N/A | N/A | N/A | N/A |
| MT | Medial terminal nucleus of the accessory optic tract | 8.46E-04 | 7.24E-04 | 5.31E-04 | 3.46E-04 | N/A | N/A | N/A | N/A |
| SNc | Substantia nigra, compact part | 5.86E-05 | 4.82E-05 | 6.02E-04 | 3.27E-04 | N/A | N/A | N/A | N/A |
| PPN | Pedunculopontine nucleus | 1.20E-06 | 1.06E-06 | 2.25E-05 | 1.51E-05 | N/A | N/A | N/A | N/A |
| NLL | Nucleus of the lateral lemniscus | 2.39E-05 | 2.62E-05 | 1.35E-04 | 6.51E-05 | N/A | N/A | N/A | N/A |
| PSV | Principal sensory nucleus of the trigeminal | 2.04E-06 | 2.47E-06 | 3.43E-07 | 3.77E-07 | N/A | N/A | N/A | N/A |
| PB | Parabrachial nucleus | 3.62E-12 | 4.44E-12 | 2.39E-05 | 2.66E-05 | N/A | N/A | N/A | N/A |
| SOC | Superior olivary complex | 2.99E-06 | 3.66E-06 | 2.45E-07 | 1.95E-07 | N/A | N/A | N/A | N/A |
| B | Barrington's nucleus | 0.00E+00 | 0.00E+00 | 7.88E-05 | 9.10E-05 | N/A | N/A | N/A | N/A |
| DTN | Dorsal tegmental nucleus | 0.00E+00 | 0.00E+00 | 0.00E+00 | 0.00E+00 | N/A | N/A | N/A | N/A |
| PDTg | Posterodorsal tegmental nucleus | 0.00E+00 | 0.00E+00 | 2.49E-06 | 2.88E-06 | N/A | N/A | N/A | N/A |
| PCG | Pontine central gray | 5.70E-06 | 4.84E-06 | 1.97E-06 | 2.27E-06 | N/A | N/A | N/A | N/A |
| PG | Pontine gray | 3.01E-03 | 3.04E-03 | 2.11E-04 | 6.69E-05 | N/A | N/A | N/A | N/A |
| PRNc | Pontine reticular nucleus, caudal part | 7.35E-07 | 5.62E-07 | 3.10E-06 | 3.32E-06 | N/A | N/A | N/A | N/A |
| SG | Supragenual nucleus | 0.00E+00 | 0.00E+00 | 0.00E+00 | 0.00E+00 | N/A | N/A | N/A | N/A |
| SUT | Supratrigeminal nucleus | 0.00E+00 | 0.00E+00 | 1.13E-06 | 1.31E-06 | N/A | N/A | N/A | N/A |
| TRN | Tegmental reticular nucleus | 5.19E-04 | 4.28E-04 | 9.37E-04 | 6.29E-04 | N/A | N/A | N/A | N/A |
| V | Motor nucleus of trigeminal | 0.00E+00 | 0.00E+00 | 9.27E-07 | 1.07E-06 | N/A | N/A | N/A | N/A |
| P5 | Peritrigeminal zone | 5.25E-08 | 6.44E-08 | 0.00E+00 | 0.00E+00 | N/A | N/A | N/A | N/A |
| Acs5 | Accessory trigeminal nucleus | 0.00E+00 | 0.00E+00 | 0.00E+00 | 0.00E+00 | N/A | N/A | N/A | N/A |
| PC5 | Parvicellular motor 5 nucleus | 9.09E-07 | 1.06E-06 | 7.07E-07 | 8.16E-07 | N/A | N/A | N/A | N/A |
| I5 | Intertrigeminal nucleus | 0.00E+00 | 0.00E+00 | 2.50E-07 | 2.89E-07 | N/A | N/A | N/A | N/A |
| CS | Superior central nucleus raphe | 1.08E-04 | 9.96E-05 | 2.96E-05 | 3.41E-05 | N/A | N/A | N/A | N/A |
| LC | Locus ceruleus | 0.00E+00 | 0.00E+00 | 4.88E-05 | 5.63E-05 | N/A | N/A | N/A | N/A |
| LDT | Laterodorsal tegmental nucleus | 4.89E-05 | 5.63E-05 | 7.85E-08 | 9.06E-08 | N/A | N/A | N/A | N/A |
| NI | Nucleus incertus | 0.00E+00 | 0.00E+00 | 0.00E+00 | 0.00E+00 | N/A | N/A | N/A | N/A |
| PRNr | Pontine reticular nucleus | 3.95E-05 | 1.20E-05 | 5.99E-05 | 6.18E-05 | N/A | N/A | N/A | N/A |
| RPO | Nucleus raphe pontis | 3.15E-06 | 2.83E-06 | 0.00E+00 | 0.00E+00 | N/A | N/A | N/A | N/A |
| SLC | Subceruleus nucleus | 1.48E-06 | 1.81E-06 | 0.00E+00 | 0.00E+00 | N/A | N/A | N/A | N/A |
| SLD | Sublaterodorsal nucleus | 0.00E+00 | 0.00E+00 | 0.00E+00 | 0.00E+00 | N/A | N/A | N/A | N/A |
| AP | Area postrema | 5.09E-07 | 6.24E-07 | 3.32E-06 | 3.83E-06 | N/A | N/A | N/A | N/A |
| ECU | External cuneate nucleus | 2.63E-06 | 2.81E-06 | 7.12E-06 | 6.04E-06 | N/A | N/A | N/A | N/A |
| NTB | Nucleus of the trapezoid body | 7.51E-06 | 9.19E-06 | 0.00E+00 | 0.00E+00 | N/A | N/A | N/A | N/A |
| NTS | Nucleus of the solitary tract | 0.00E+00 | 0.00E+00 | 1.02E-06 | 9.36E-07 | N/A | N/A | N/A | N/A |
| SPVC | Spinal nucleus of the trigeminal, caudal part | 2.36E-06 | 2.88E-06 | 3.07E-07 | 3.54E-07 | N/A | N/A | N/A | N/A |
| SPVI | Spinal nucleus of the trigeminal, interpolar part | 4.10E-07 | 5.02E-07 | 4.31E-07 | 3.73E-07 | N/A | N/A | N/A | N/A |
| SPVO | Spinal nucleus of the trigeminal, oral part | 2.55E-06 | 3.13E-06 | 1.73E-07 | 2.00E-07 | N/A | N/A | N/A | N/A |
| Pa5 | Paratrigeminal nucleus | 6.49E-05 | 7.95E-05 | 4.26E-06 | 4.81E-06 | N/A | N/A | N/A | N/A |
| VI | Abducens nucleus | 0.00E+00 | 0.00E+00 | 0.00E+00 | 0.00E+00 | N/A | N/A | N/A | N/A |
| VII | Facial motor nucleus | 2.28E-06 | 2.75E-06 | 1.73E-07 | 1.26E-07 | N/A | N/A | N/A | N/A |
| ACVII | Accessory facial motor nucleus | 0.00E+00 | 0.00E+00 | 0.00E+00 | 0.00E+00 | N/A | N/A | N/A | N/A |
| AMB | Nucleus ambiguus | 8.40E-09 | 1.03E-08 | 8.12E-09 | 9.38E-09 | N/A | N/A | N/A | N/A |
| DMX | Dorsal motor nucleus of the vagus nerve | 0.00E+00 | 0.00E+00 | 0.00E+00 | 0.00E+00 | N/A | N/A | N/A | N/A |
| GRN | Gigantocellular reticular nucleus | 7.07E-06 | 7.37E-06 | 2.82E-06 | 2.84E-06 | N/A | N/A | N/A | N/A |
| ICB | Infracerebellar nucleus | 0.00E+00 | 0.00E+00 | 0.00E+00 | 0.00E+00 | N/A | N/A | N/A | N/A |
| IO | Inferior olivary complex | 8.35E-06 | 1.02E-05 | 1.69E-05 | 1.78E-05 | N/A | N/A | N/A | N/A |
| IRN | Intermediate reticular nucleus | 4.43E-06 | 5.21E-06 | 1.48E-06 | 1.71E-06 | N/A | N/A | N/A | N/A |
| ISN | Inferior salivatory nucleus | 0.00E+00 | 0.00E+00 | 0.00E+00 | 0.00E+00 | N/A | N/A | N/A | N/A |
| LIN | Linear nucleus of the medulla | 0.00E+00 | 0.00E+00 | 5.21E-06 | 6.02E-06 | N/A | N/A | N/A | N/A |
| LRN | Lateral reticular nucleus | 3.35E-05 | 3.91E-05 | 2.01E-06 | 2.11E-06 | N/A | N/A | N/A | N/A |
| MARN | Magnocellular reticular nucleus | 4.02E-06 | 4.55E-06 | 5.23E-05 | 5.89E-05 | N/A | N/A | N/A | N/A |
| MDRN | Medullary reticular nucleus | 2.82E-05 | 3.45E-05 | 6.40E-07 | 4.44E-07 | N/A | N/A | N/A | N/A |
| PARN | Parvicellular reticular nucleus | 3.70E-06 | 4.44E-06 | 1.88E-06 | 1.35E-06 | N/A | N/A | N/A | N/A |
| PAS | Parasolitary nucleus | 0.00E+00 | 0.00E+00 | 0.00E+00 | 0.00E+00 | N/A | N/A | N/A | N/A |
| PPY | Parapyramidal nucleus | 1.50E-05 | 1.83E-05 | 0.00E+00 | 0.00E+00 | N/A | N/A | N/A | N/A |
| x | Nucleus x | 0.00E+00 | 0.00E+00 | 0.00E+00 | 0.00E+00 | N/A | N/A | N/A | N/A |
| XII | Hypoglossal nucleus | 0.00E+00 | 0.00E+00 | 1.40E-06 | 1.61E-06 | N/A | N/A | N/A | N/A |
| y | Nucleus y | 0.00E+00 | 0.00E+00 | 0.00E+00 | 0.00E+00 | N/A | N/A | N/A | N/A |
| RM | Nucleus raphe magnus | 7.93E-06 | 9.71E-06 | 1.50E-06 | 1.73E-06 | N/A | N/A | N/A | N/A |
| RPA | Nucleus raphe pallidus | 2.81E-04 | 3.17E-04 | 1.27E-04 | 1.47E-04 | N/A | N/A | N/A | N/A |
| RO | Nucleus raphe obscurus | 2.98E-06 | 3.64E-06 | 0.00E+00 | 0.00E+00 | N/A | N/A | N/A | N/A |
| FN | Fastigial nucleus | 1.56E-06 | 1.92E-06 | 4.98E-07 | 3.47E-07 | N/A | N/A | N/A | N/A |
| IP | Interposed nucleus | 0.00E+00 | 0.00E+00 | 1.24E-07 | 1.43E-07 | N/A | N/A | N/A | N/A |
| DN | Dentate nucleus | 0.00E+00 | 0.00E+00 | 2.10E-06 | 1.42E-06 | N/A | N/A | N/A | N/A |
| VeCB | Vestibulocerebellar nucleus | 1.31E-07 | 1.60E-07 | 1.01E-06 | 1.17E-06 | N/A | N/A | N/A | N/A |

**Table S2.** Comparison with previous whole-brain characterization of input and/or output of specific BF subnuclei. For reference, our AP coordinates for HDB and SI were 0.6 and -0.23, respectively. Note that Chen et al., 2023 was done on the Nucleus Basalis Magnocellularis (NBM), so here we compare it to our SI data due to their proximity.

| **BF subnucleus** | **Paper** | **Input** | **Output** |
|---|---|---|---|
| HDB (AP Bregma 0.1 mm) | Do et al. 2016 | *Compatible*: major input from the piriform, prefrontal, orbitofrontal, and insular cortices.<br>*Discrepant*: unlike them, we did not find substantial input from the central amygdala.<br>*Unclear*: they found major input from the caudate putamen and the lateral hypothalamus, but these were mostly masked here, due to proximity to the injection site. | *Compatible*: major output to hippocampal regions, frontal and posterior cortex.<br>*Discrepant*: unlike them, we identified relatively fewer projections to the piriform, motor and somatosensory cortices. |
| HDB (AP Bregma -0.1 mm) | Hu et al. 2016 | *Compatible*: major input from the piriform, prefrontal, orbitofrontal, and insular cortices.<br>*Discrepant*: unlike them, we did not find substantial input from the central amygdala.<br>*Unclear*: they found major input from the caudate putamen and the lateral hypothalamus, but these were mostly masked here, due to proximity to the injection site. | N/A |
| NBM (AP Bregma -0.7 mm)<br>(i.e., closer to our SI than our HDB coordinate) | Chen et al. 2023 | *Compatible*: major input from the central amygdala, and insular and motor cortices.<br>*Discrepant*: unlike them, we identified more input from the piriform and prefrontal cortices to the SI.<br>*Unclear*: they found major input from the caudate putamen, but these were mostly masked here, due to proximity to the injection site. | *Compatible*: major output to the anterior motor and somatosensory cortices and no projections to the hippocampus. |
| MS-HDB (AP Bregma +0.1 mm) | Chen et al. 2026 | N/A | *Compatible*: major output to hippocampal regions as well as frontal and posterior cortices.<br>*Discrepant*: unlike them, we identified relatively fewer projections to the basolateral amygdala and piriform cortex. |

## References

1. Zaborszky, L., van den Pol, A., and Gyengési, E. (2011). The Basal Forebrain Cholinergic Projection System in Mice. In The mouse nervous system, C. Watson, G. Paxinos, and L. Puelles, eds. (Elsevier), pp. 684–709.

2. Rye, D.B., Wainer, B.H., Mesulam, M.M., Mufson, E.J., and Saper, C.B. (1984). Cortical projections arising from the basal forebrain: a study of cholinergic and noncholinergic components employing combined retrograde tracing and immunohistochemical localization of choline acetyltransferase. Neuroscience 13, 627–643.

3. Sarter, M., and Bruno, J.P. (2002). The neglected constituent of the basal forebrain corticopetal projection system: GABAergic projections. Eur. J. Neurosci. 15, 1867–1873.

4. Lee, S.-H., and Dan, Y. (2012). Neuromodulation of brain states. Neuron 76, 209–222.

5. Buzsaki, G., Bickford, R.G., Ponomareff, G., Thal, L.J., Mandel, R., and Gage, F.H. (1988). Nucleus basalis and thalamic control of neocortical activity in the freely moving rat. J. Neurosci. 8, 4007–4026.

6. Lohani, S., Moberly, A.H., Benisty, H., Landa, B., Jing, M., Li, Y., Higley, M.J., and Cardin, J.A. (2022). Spatiotemporally heterogeneous coordination of cholinergic and neocortical activity. Nat. Neurosci. 25, 1706–1713.

7. Neyhart, E., Zhou, N., Munn, B.R., Law, R.G., Smith, C., Mridha, Z.H., Blanco, F.A., Li, G., Li, Y., Hu, M., et al. (2024). Cortical acetylcholine dynamics are predicted by cholinergic axon activity and behavior state. Cell Rep. 43, 114808.

8. Collins, L., Francis, J., Emanuel, B., and McCormick, D.A. (2023). Cholinergic and noradrenergic axonal activity contains a behavioral-state signal that is coordinated across the dorsal cortex. Elife 12. 10.7554/eLife.81826.

9. Fu, Y., Tucciarone, J.M., Espinosa, J.S., Sheng, N., Darcy, D.P., Nicoll, R.A., Huang, Z.J., and Stryker, M.P. (2014). A Cortical Circuit for Gain Control by Behavioral State. Cell 156, 1139–1152.

10. Lee, A.M., Hoy, J.L., Bonci, A., Wilbrecht, L., Stryker, M.P., and Niell, C.M. (2014). Identification of a brainstem circuit regulating visual cortical state in parallel with locomotion. Neuron 83, 455–466.

11. Harrison, T.C., Pinto, L., Brock, J.R., and Dan, Y. (2016). Calcium imaging of basal forebrain activity during innate and learned behaviors. Front. Neural Circuits 10, 36.

12. Zhu, F., Elnozahy, S., Lawlor, J., and Kuchibhotla, K.V. (2023). The cholinergic basal forebrain provides a parallel channel for state-dependent sensory signaling to auditory cortex. Nat. Neurosci. 26, 810–819.

13. Froemke, R.C., Merzenich, M.M., and Schreiner, C.E. (2007). A synaptic memory trace for cortical receptive field plasticity. Nature 450, 425–429.

14. Kilgard, M.P., and Merzenich, M.M. (1998). Cortical map reorganization enabled by nucleus basalis activity. Science 279, 1714–1718.

15. Rajebhosale, P., Ananth, M.R., Kim, R., Crouse, R., Jiang, L., López-Hernández, G., Zhong, C., Arty, C., Wang, S., Jone, A., et al. (2024). Functionally refined encoding of threat memory by distinct populations of basal forebrain cholinergic projection neurons. Elife 13, e86581.

16. Hasselmo, M.E. (2006). The role of acetylcholine in learning and memory. Curr. Opin. Neurobiol. 16, 710–715.

17. Chen, N., Sugihara, H., Sharma, J., Perea, G., Petravicz, J., Le, C., and Sur, M. (2012). Nucleus basalis-enabled stimulus-specific plasticity in the visual cortex is mediated by astrocytes. Proceedings of the National Academy of Sciences 109, E2832–E2841.

18. Hangya, B., Ranade, S.P., Lorenc, M., and Kepecs, A. (2015). Central cholinergic neurons are rapidly recruited by reinforcement feedback. Cell 162, 1155–1168.

19. Robert, B., Kimchi, E.Y., Watanabe, Y., Chakoma, T., Jing, M., Li, Y., and Polley, D.B. (2021). A functional topography within the cholinergic basal forebrain for encoding sensory cues and behavioral reinforcement outcomes. Elife 10. 10.7554/eLife.69514.

20. Crouse, R.B., Kim, K., Batchelor, H.M., Girardi, E.M., Kamaletdinova, R., Chan, J., Rajebhosale, P., Pittenger, S.T., Role, L.W., Talmage, D.A., et al. (2020). Acetylcholine is released in the basolateral amygdala in response to predictors of reward and enhances the learning of cue-reward contingency. Elife 9, e57335.

21. Chubykin, A.A., Roach, E.B., Bear, M.F., and Shuler, M.G.H. (2013). A cholinergic mechanism for reward timing within primary visual cortex. Neuron 77, 723–735.

22. Laszlovszky, T., Schlingloff, D., Hegedüs, P., Freund, T.F., Gulyás, A., Kepecs, A., and Hangya, B. (2020). Distinct synchronization, cortical coupling and behavioral function of two basal forebrain cholinergic neuron types. Nat. Neurosci. 23, 992–1003.

23. Herrero, J.L., Roberts, M.J., Delicato, L.S., Gieselmann, M.A., Dayan, P., and Thiele, A. (2008). Acetylcholine contributes through muscarinic receptors to attentional modulation in V1. Nature 454, 1110–1114.

24. Pinto, L., Goard, M.J., Estandian, D., Xu, M., Kwan, A.C., Lee, S.-H., Harrison, T.C., Feng, G., and Dan, Y. (2013). Fast modulation of visual perception by basal forebrain cholinergic neurons. Nat. Neurosci. 16, 1857–1863.

25. Parikh, V., Kozak, R., Martinez, V., and Sarter, M. (2007). Prefrontal acetylcholine release controls cue detection on multiple timescales. Neuron 56, 141–154.

26. Yu, B., Yue, Y., Ren, C., Yun, R., Lim, B., and Komiyama, T. (2025). Cholinergic feedback for modality-and context-specific modulation of sensory representations. Science 388, 1324–1329.

27. Luo, J.K., Tan, J., Myhre, E., Salvino, P., Hark, A., and Pinto, L. (2025). A cholinergic mechanism orchestrating task-dependent computation across the cortex. bioRxivorg. 10.1101/2025.11.26.690825.

28. Zou, J., Willem de Gee, J., Mridha, Z., Trinh, S., Erskine, A., Jing, M., Yao, J., Walker, S., Li, Y., McGinley, M., et al. (2024). Goal-directed motor actions drive acetylcholine dynamics in sensory cortex. eLife. 10.7554/elife.96931.1.

29. Minces, V., Pinto, L., Dan, Y., and Chiba, A.A. (2017). Cholinergic shaping of neural correlations. Proc. Natl. Acad. Sci. U. S. A. 114, 5725–5730.

30. Galvin, V.C., and Disney, A.A. (2026). Perceptual performance and V1 neural activity are exquisitely sensitive to small changes in thalamocortical gain. bioRxivorg, 2026.02.22.707315. 10.64898/2026.02.22.707315.

31. Khalighinejad, N., Bongioanni, A., Verhagen, L., Folloni, D., Attali, D., Aubry, J.-F., Sallet, J., and Rushworth, M.F.S. (2020). A basal forebrain-cingulate circuit in macaques decides it is time to act. Neuron 105, 370–384.e8.

32. Wang, H., Ortega, H.K., Kelly, E.B., Indajang, J., Savalia, N.K., Glaeser-Khan, S., Feng, J., Li, Y., Kaye, A.P., and Kwan, A.C. (2025). Frontal noradrenergic and cholinergic transients exhibit distinct spatiotemporal dynamics during competitive decision-making. Sci. Adv. 11, eadr9916.

33. Garg, R., Qiu, Q., and Yu, C.R. (2025). Basal forebrain cholinergic input mediates adaptive attention allocation to enhance olfactory discrimination. PLoS Biol. 23, e3003374.

34. Bigl, V., Woolf, N.J., and Butcher, L.L. (1982). Cholinergic projections from the basal forebrain to frontal, parietal, temporal, occipital, and cingulate cortices: a combined fluorescent tracer and acetylcholinesterase analysis. Brain Res. Bull. 8, 727–749.

35. Saper, C.B. (1984). Organization of cerebral cortical afferent systems in the rat. II. Magnocellular basal nucleus. J. Comp. Neurol. 222, 313–342.

36. Gielow, M.R., and Zaborszky, L. (2017). The Input-Output Relationship of the Cholinergic Basal Forebrain. Cell Rep. 18, 1817–1830.

37. Chen, Z., Liu, Y., Yang, Y., Wang, L., Qin, M., Jiang, Z., Xu, M., and Zhang, S. (2025). Whole-brain mapping of basal forebrain cholinergic neurons reveals a long-range reciprocal input-output loop between distinct subtypes. Sci. Adv. 10.1126/sciadv.adt1617.

38. Muñoz, W., and Rudy, B. (2014). Spatiotemporal specificity in cholinergic control of neocortical function. Curr. Opin. Neurobiol. 26, 149–160.

39. Zaborszky, L., Csordas, A., Mosca, K., Kim, J., Gielow, M.R., Vadasz, C., and Nadasdy, Z. (2015). Neurons in the basal forebrain project to the cortex in a complex topographic organization that reflects corticocortical connectivity patterns: an experimental study based on retrograde tracing and 3D reconstruction. Cereb. Cortex 25, 118–137.

40. Záborszky, L., Gombkoto, P., Varsanyi, P., Gielow, M.R., Poe, G., Role, L.W., Ananth, M., Rajebhosale, P., Talmage, D.A., Hasselmo, M.E., et al. (2018). Specific basal forebrain-cortical cholinergic circuits coordinate cognitive operations. J. Neurosci. 38, 9446–9458.

41. Varsanyi, P., Alloway, K., Chavez, C., Gielow, M.R., Gombkoto, P., Kondo, H., Nadasdy, Z., and Zaborszky, L. (2025). Hierarchical organization of the forebrain cholinergic system in rats. iScience 28, 112001.

42. Eckenstein, F.P., Baughman, R.W., and Quinn, J. (1988). An anatomical study of cholinergic innervation in rat cerebral cortex. Neuroscience 25, 457–474.

43. Carnes, K.M., Fuller, T.A., and Price, J.L. (1990). Sources of presumptive glutamatergic/aspartatergic afferents to the magnocellular basal forebrain in the rat. J. Comp. Neurol. 302, 824–852.

44. Gaykema, R.P., Luiten, P.G., Nyakas, C., and Traber, J. (1990). Cortical projection patterns of the medial septum-diagonal band complex. J. Comp. Neurol. 293, 103–124.

45. Gaykema, R.P., van Weeghel, R., Hersh, L.B., and Luiten, P.G. (1991). Prefrontal cortical projections to the cholinergic neurons in the basal forebrain. J. Comp. Neurol. 303, 563–583.

46. Zaborszky, L., Gaykema, R.P., Swanson, D.J., and Cullinan, W.E. (1997). Cortical input to the basal forebrain. Neuroscience 79, 1051–1078.

47. Mesulam, M.M., Mufson, E.J., Wainer, B.H., and Levey, A.I. (1983). Central cholinergic pathways in the rat: an overview based on an alternative nomenclature (Ch1-Ch6). Neuroscience 10, 1185– 1201.

48. Mesulam, M.M., Mufson, E.J., Levey, A.I., and Wainer, B.H. (1983). Cholinergic innervation of cortex by the basal forebrain: cytochemistry and cortical connections of the septal area, diagonal band nuclei, nucleus basalis (substantia innominata), and hypothalamus in the rhesus monkey. J. Comp. Neurol. 214, 170–197.

49. Laplante, F., Morin, Y., Quirion, R., and Vaucher, E. (2005). Acetylcholine release is elicited in the visual cortex, but not in the prefrontal cortex, by patterned visual stimulation: a dual in vivo microdialysis study with functional correlates in the rat brain. Neuroscience 132, 501–510.

50. Li, X., Yu, B., Sun, Q., Zhang, Y., Ren, M., Zhang, X., Li, A., Yuan, J., Madisen, L., Luo, Q., et al. (2018). Generation of a whole-brain atlas for the cholinergic system and mesoscopic projectome analysis of basal forebrain cholinergic neurons. Proc. Natl. Acad. Sci. U. S. A. 115, 415–420.

51. Wu, H., Williams, J., and Nathans, J. (2014). Complete morphologies of basal forebrain cholinergic neurons in the mouse. Elife 3, e02444.

52. Do, J.P., Xu, M., Lee, S.-H., Chang, W.-C., Zhang, S., Chung, S., Yung, T.J., Fan, J.L., Miyamichi, K., Luo, L., et al. (2016). Cell type-specific long-range connections of basal forebrain circuit. Elife 5. 10.7554/eLife.13214.

53. Chen, Y., Tian, J., Luo, S., Ren, M., Jiang, T., Gong, H., Luo, Q., Li, A., and Li, X. (2026). Synaptic-level organization of long-range cholinergic circuits in the basal forebrain. Commun. Biol., 1–11.

54. Hu, R., Jin, S., He, X., Xu, F., and Hu, J. (2016). Whole-brain monosynaptic afferent inputs to basal forebrain cholinergic system. Front. Neuroanat. 10, 98.

55. Chen, Z.-Y., Yang, Y.-L., Li, M., Gao, L., Qu, W.-M., Huang, Z.-L., and Yuan, X.-S. (2023). Whole-brain neural connectivity to cholinergic neurons in the nucleus basalis of Meynert. J. Neurochem. 166, 233–247.

56. Oh, S.W., Harris, J.A., Ng, L., Winslow, B., Cain, N., Mihalas, S., Wang, Q., Lau, C., Kuan, L., Henry, A.M., et al. (2014). A mesoscale connectome of the mouse brain. Nature 508, 207–214.

57. Kuan, L., Li, Y., Lau, C., Feng, D., Bernard, A., Sunkin, S.M., Zeng, H., Dang, C., Hawrylycz, M., and Ng, L. (2015). Neuroinformatics of the Allen Mouse Brain Connectivity Atlas. Methods 73, 4–17.

58. Wang, Q., Ding, S.-L., Li, Y., Royall, J., Feng, D., Lesnar, P., Graddis, N., Naeemi, M., Facer, B., Ho, A., et al. (2020). The Allen Mouse Brain Common Coordinate Framework: A 3D reference atlas. Cell 181, 936–953.e20.

59. Allaway, K.C., Muñoz, W., Tremblay, R., Sherer, M., Herron, J., Rudy, B., Machold, R., and Fishell, G. (2020). Cellular birthdate predicts laminar and regional cholinergic projection topography in the forebrain. Elife 9, e63249.

60. Yates, S.C., Groeneboom, N.E., Coello, C., Lichtenthaler, S.F., Kuhn, P.-H., Demuth, H.-U., Hartlage-Rübsamen, M., Roßner, S., Leergaard, T., Kreshuk, A., et al. (2019). QUINT: Workflow for quantification and spatial analysis of features in histological images from rodent brain. Front. Neuroinform. 13. 10.3389/fninf.2019.00075.

61. Henny, P., and Jones, B.E. (2008). Projections from basal forebrain to prefrontal cortex comprise cholinergic, GABAergic and glutamatergic inputs to pyramidal cells or interneurons. Eur. J. Neurosci. 27, 654–670.

62. Golmayo, L., Nuñez, A., and Zaborszky, L. (2003). Electrophysiological evidence for the existence of a posterior cortical-prefrontal-basal forebrain circuitry in modulating sensory responses in visual and somatosensory rat cortical areas. Neuroscience 119, 597–609.

63. Nelson, C.L., Sarter, M., and Bruno, J.P. (2005). Prefrontal cortical modulation of acetylcholine release in posterior parietal cortex. Neuroscience 132, 347–359.

64. Rasmusson, D.D., Smith, S.A., and Semba, K. (2007). Inactivation of prefrontal cortex abolishes cortical acetylcholine release evoked by sensory or sensory pathway stimulation in the rat. Neuroscience 149, 232–241.

65. Biderman, D., Whiteway, M.R., Hurwitz, C., Greenspan, N., Lee, R.S., Vishnubhotla, A., Warren, R., Pedraja, F., Noone, D., Schartner, M.M., et al. (2024). Lightning Pose: improved animal pose estimation via semi-supervised learning, Bayesian ensembling and cloud-native open-source tools. Nat. Methods 21, 1316–1328.

66. Xu, M., Chung, S., Zhang, S., Zhong, P., Ma, C., Chang, W.-C., Weissbourd, B., Sakai, N., Luo, L., Nishino, S., et al. (2015). Basal forebrain circuit for sleep-wake control. Nat. Neurosci., 1–9.

67. Zaborszky, L., and Duque, A. (2000). Local synaptic connections of basal forebrain neurons. Behav. Brain Res. 115, 143–158.

68. Alitto, H.J., and Dan, Y. (2012). Cell-type-specific modulation of neocortical activity by basal forebrain input. Front. Syst. Neurosci. 6, 79.

69. McCormick, D.A., and Prince, D.A. (1985). Two types of muscarinic response to acetylcholine in mammalian cortical neurons. Proc. Natl. Acad. Sci. U. S. A. 82, 6344–6348.

70. Picciotto, M.R., Higley, M.J., and Mineur, Y.S. (2012). Acetylcholine as a Neuromodulator: Cholinergic Signaling Shapes Nervous System Function and Behavior. Neuron 76, 116–129.

71. Pinto, L., and Dan, Y. (2015). Cell-Type-Specific Activity in Prefrontal Cortex during Goal-Directed Behavior. Neuron 87, 437–450.

72. Le Merre, P., Heining, K., Slashcheva, M., Jung, F., Moysiadou, E., Guyon, N., Yahya, R., Park, H., Wernstal, F., and Carlén, M. (2026). A prefrontal cortex map based on single-neuron activity. Nat. Neurosci., 1–9.

73. Jung, M.W., Qin, Y., Mcnaughton, B.L., and Barnes, C.A. (1998). Firing characteristics of deep layer neurons in prefrontal cortex in rats performing spatial working memory tasks. Cereb. Cortex 8, 437–450.

74. Rodgers, C.C., and DeWeese, M.R. (2014). Neural Correlates of Task Switching in Prefrontal Cortex and Primary Auditory Cortex in a Novel Stimulus Selection Task for Rodents. Neuron 82, 1157–1170.

75. Cole, N., Harvey, M., Myers-Joseph, D., Gilra, A., and Khan, A.G. (2024). Prediction-error signals in anterior cingulate cortex drive task-switching. Nat. Commun. 15, 1–20.

76. Li, S.-J., Hangya, B., Gupta, U., Singh, H., Fischer, K.B., Sturgill, J.F., Callaway, E.M., and Kepecs, A. (2026). A disinhibitory basal forebrain-to-cortex projection supports sustained attention. Cell 189, 4140–4158.e9.

77. Guo, Z.V., Li, N., Huber, D., Ophir, E., Gutnisky, D., Ting, J.T., Feng, G., and Svoboda, K. (2014). Flow of cortical activity underlying a tactile decision in mice. Neuron 81, 179–194.

78. Costa, R.M., Luo, J.K., Salvino, P.S., Ackert-Smith, L.A., Ibarra, S.M., and Pinto, L. (2025). Cognitive processes are disentangled at cortex-wide scales. bioRxivorg. 10.1101/2025.07.24.666672.

79. Groeneboom, N.E., Yates, S.C., Puchades, M.A., and Bjaalie, J.G. (2020). Nutil: A pre- and post-processing toolbox for histological rodent brain section images. Front. Neuroinform. 14, 37.

80. Carey, H., Pegios, M., Martin, L., Saleeba, C., Turner, A.J., Everett, N.A., Bjerke, I.E., Puchades, M.A., Bjaalie, J.G., and McMullan, S. (2023). DeepSlice: rapid fully automatic registration of mouse brain imaging to a volumetric atlas. Nat. Commun. 14, 5884.

81. Puchades, M.A., Csucs, G., Ledergerber, D., Leergaard, T.B., and Bjaalie, J.G. (2019). Spatial registration of serial microscopic brain images to three-dimensional reference atlases with the QuickNII tool. PLoS One 14, e0216796.

82. Gurdon, B., Yates, S.C., Csucs, G., Groeneboom, N.E., Hadad, N., Telpoukhovskaia, M., Ouellette, A., Ouellette, T., O’Connell, K.M.S., Singh, S., et al. (2024). Detecting the effect of genetic diversity on brain composition in an Alzheimer’s disease mouse model. Commun. Biol. 7. 10.1038/s42003-024-06242-1.

83. Berg, S., Kutra, D., Kroeger, T., Straehle, C.N., Kausler, B.X., Haubold, C., Schiegg, M., Ales, J., Beier, T., Rudy, M., et al. (2019). Ilastik: Interactive machine learning for (bio)image analysis. Nat. Methods 16, 1226–1232.

84. Ma, Y., Shaik, M.A., Kim, S.H., Kozberg, M.G., Thibodeaux, D.N., Zhao, H.T., Yu, H., and Hillman, E.M.C. (2016). Wide-field optical mapping of neural activity and brain haemodynamics: considerations and novel approaches. Philos. Trans. R. Soc. Lond. B Biol. Sci. 371, 20150360.

